# Morphotype-Resolved 3D Morphometry Reveals a Structure–Density–Location Coupling in Mitochondrial Networks

**DOI:** 10.64898/2026.03.19.712811

**Authors:** Anshu Singh, Arun Yadav, Aneesh Deshmukh, Arjun Singh, Riva Verma, Kate White, Jitin Singla

## Abstract

Mitochondrial architecture plays a critical role in cellular function, yet how organelle structure, metabolic density, and subcellular position are jointly remodeled across whole cells remains poorly understood. We applied quantitative 3D soft X-ray tomography to analyze intact INS-1E cells in a native, cryo-hydrated state. By integrating morphometric profiling with voxel-level linear absorption coefficients (LAC) and contour-based radial mapping, we tracked the structural, biochemical, and spatial remodeling of fragmented, intermediate, and interconnected mitochondrial morphotypes under high glucose and Exendin-4 stimulation. High glucose induces morphotype-specific hypertrophy, fission, and perinuclear redistribution of low-density fragments. Co-stimulation with Exendin-4 stabilizes interconnected networks and increases metabolic density at the cell periphery. Morphotype-resolved analysis uncovers a structure–density–location coupling in which mitochondrial shape, macromolecular packing, and radial position shift in concert. These results offer a quantitative framework and high-resolution spatial constraints for whole-cell modeling of organelle dynamics.

**Highlights:**

- Mitochondrial morphotypes occupy distinct regions during metabolic stimulation
- High glucose drives early mitochondrial swelling, fission, and spatial dispersion
- Exendin-4 prevents structural collapse and preserves dense interconnected networks
- Morphotype-resolved analysis uncovers a structure–density–location coupling in mitochondrial networks.

**Graphical Abstract:** 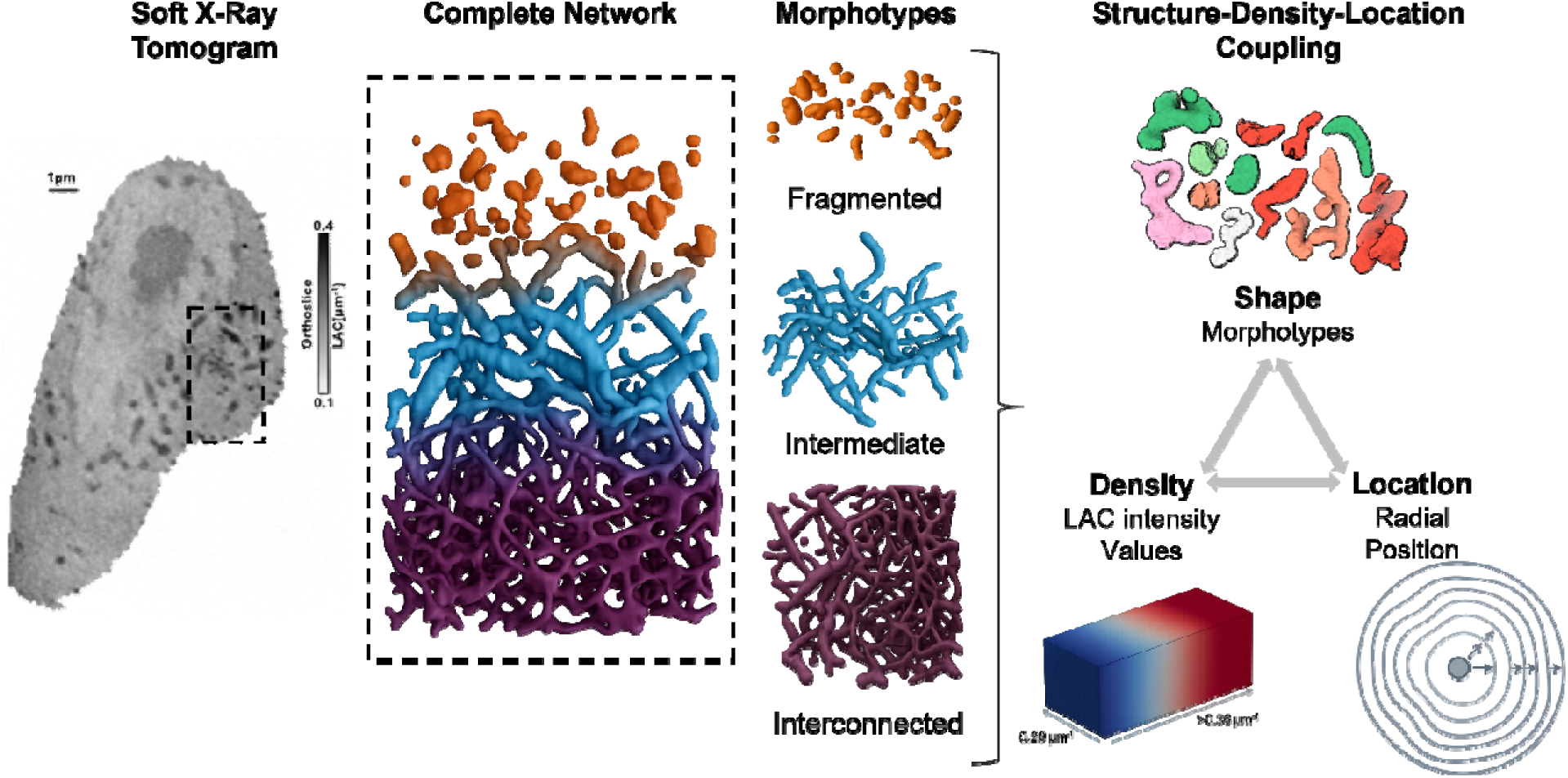

## INTRODUCTION

Mitochondria are highly dynamic organelles that constantly reshape their morphology through fusion and fission to maintain energy production and cellular homeostasis^1,2^. During fission, structural proteins and the endoplasmic reticulum help constrict the outer membrane, while energy-driven proteins assemble around the mitochondrion to tighten and complete the split^3,4^. In contrast, fusion involves membrane-associated proteins that bring mitochondria together, merge membranes, and mix contents such as DNA and proteins to sustain functional stability^5–7^. When this dynamic balance is disturbed, it can lead to mitochondrial dysfunction and disease, whereas maintaining balanced dynamics promotes metabolic health and longevity^7^. As structure-function is closely tied, not just for proteins, but for larger biological systems as well, mitochondrial structural changes have been observed to correlate with mitochondrial dysfunction^8,9^. For instance, excessive fragmentation, as seen in ischemia, can cause cell death in heart tissue^10^. Similarly, excessive mitochondrial fragmentation in pancreatic β-cell as observed under high glucose conditions, is associated with reduced ATP synthesis and impaired glucose-stimulated insulin secretion^11,12^. Fragmented mitochondria in diabetic β-cell models show reduced oxidative phosphorylation capacity along with weakened metabolic responses to glucose stimulation^13^. A similar trend is also observed when mitochondrial fusion proteins are pharmacologically inhibited, leading to fragmented mitochondrial morphologies accompanied by a compromised capacity of insulin secretion^14^. Further, abnormal mitochondrial division is also reported to contribute to insulin resistance^15^. **Given this tight structure-function coupling, mitochondrial architecture serves as a critical biomarker for cellular metabolic state and health.** As such, identifying and measuring morphological parameters accurately will be critical for the characterization of the structural state of mitochondria, as well as for targeting mitochondrial morphology for therapeutic intervention^16^. It will also facilitate the modelling of mitochondrial dynamics at the level of the cell and across the entire cellular environment^17–21^.

Over the past decade, studies investigating mitochondria have gradually shifted from largely qualitative descriptions of organelles being either “fragmented” or “fused” to more towards quantitative morphometric characterization or profiling. Much of this transition was enabled by the advancement of high-content imaging strategies together with morphology membrane potential binning innovative approaches that allowed classification of mitochondrial phenotypes directly in the living cells^22^. During this period, there has been considerable advancement in the quantitative assessment of mitochondrial morphology, morphometric analyses can capture the functional states of organelle. Earlier high-impact works, primarily relied on 2D confocal imaging to measure qualitative or semi-quantitative descriptors of fragmentation, tubular networks, and fission–fusion imbalance^22–25^. As the field progressed, method-driven studies introduced more systematic quantitative metrics, including measurements of mitochondrial volume, area, perimeter, form factor, branch length and branch points, often extracted using tools such as ImageJ^26^. Parallelly some studies also established 3D volumetric reconstructions of mitochondrial networks, reporting additional parameters such as object count and overall network connectivity using tools like MitoGraph^27^, Imaris^28^ or manual segmentation^25,29–31^. More recent computational methods have moved toward automated extraction of 3D organelle shape and topology allowing more consistent structural characterization of mitochondrial networks^32,33^.

Despite its importance, quantitative analysis of mitochondrial morphology in intact cells has remained challenging, hindered by both imaging constraints and analytical oversimplification. Most current morphometric studies rely on fluorescence or electron microscopy; however, fluorescence is limited by photobleaching, anisotropic resolution and labeling perturbations^34,35^, and electron microscopy techniques like Transmission electron microscopy (TEM) and focused-ion-beam scanning electron microscopy (FIB-SEM) require extensive sample preparation like fixation, dehydration, and sectioning, which can obscure native cellular structures, prohibit imaging in physiological conditions and restricts large sample collection^36,37^. Furthermore, cryo-electron tomography^38^ (cryo-ET) while preserving near-native states at molecular resolution (∼30 Å), is restricted by a small field of view that precludes whole-cell analysis^39–42^. Consequently, many studies have resorted to coarse, binary classification (e.g. “fused” vs. “fragmented”) failing to capture the true morphological continuum of intermediate network states. Even where more sophisticated machine learning techniques has been applied to fluorescence data^22,43,44^, these analyses remain limited by (i) non-native imaging conditions, (ii) tool-specific parameter limitations, and (iii) a reliance on 2D projections. Crucially, a quantitative characterization of mitochondrial spatial organization at the whole-cell level remains limited and a morphotype-resolved mapping of mitochondrial intensity across a complete network has yet to be explored. These technical and methodological constraints have restricted morphometric analysis to isolated snapshots, precluding a comprehensive quantification of mitochondrial landscape in its native, 3D whole-cell context.

To address these limitations, we applied quantitative 3D soft X-ray tomography (SXT) to image intact INS-1E pancreatic β-cells in a near-native cryo-hydrated state. SXT operates within the “water window” (284–543 eV), exploiting the natural X-ray absorption of carbon-rich structures to provide high-contrast, label-free imaging of organelles at ∼30–50 nm resolution. Unlike traditional electron microscopy, SXT requires no chemical fixation or sectioning, preserving the cellular ultrastructure. Furthermore, the voxel intensity value, called the linear absorption coefficient (LAC) in SXT, is directly proportional to local biomolecular density, enabling us to quantify organelle composition alongside morphology. Previously, SXT investigation have mapped cellular ultrastructure in yeast, yeast suspension^45^, viruses^46^, mammalian cells^47^ and β-cells^48–52^. SXT studies have provided a general ultrastructural characterization of β-cell organelles, including quantitative measurements of organelle volumes and spatial distribution^50^, detailed analysis of mitochondria–vesicle contact sites^48^, and deeper insights into insulin-vesicle maturation under distinct secretory stimuli^51^. While these studies provided a ultrastructural blueprint of the β-cell, they did not specifically address the heterogeneity of the mitochondrial network. Building on these foundational observations, the present work extends this framework by introducing mitochondrial network architecture integrated with voxel-level density mapping as a quantitative descriptor of cellular metabolic state. Crucially, whether mitochondrial morphotype, biomolecular density, and subcellular position are independently regulated or shift in a coordinated manner remains unexplored.

In this study, we bridge the gap between whole-cell imaging and detailed morphological analysis. By analyzing 55 cells across a temporal gradient (1, 5, and 30 minutes) of high-glucose and Exendin-4 stimulation, we reveal how metabolic “shocks” rewire the mitochondrial network. Aligning with the previous literature^53–55^, our analysis stratifies mitochondria into three distinct morphotypes: fragmented, intermediate, and interconnected and using unsupervised k-means clustering further resolves the geometric “fingerprints” of the transitional intermediate pool. We provide a comprehensive assessment of how the β-cell mitochondrial landscape responds under therapeutic intervention, by using 3D volumetric reconstructions along with radial enrichment mapping and also integrating LAC intensity profiling in this study.

## RESULTS

### Global Mitochondrial Network Remodeling

To capture the spatiotemporal dynamics of the entire mitochondrial network during metabolic stimulation we performed a comprehensive quantitative assessment of mitochondrial morphology and biochemical density in *n* = 55 whole-cell tomograms (Figure S1). Cells were analyzed under distinct metabolic states: an unstimulated control (NS), stimulation with 25 mM high glucose, 10 nM Exendin-4 (Ex-4), or a combination of both at specific intervals of 1, 5 and 30 minutes. Quantification metrics such as volume, surface area, mitochondrial complexity, and linear absorption coefficient (LAC) of global network revealed that the mitochondrial network undergoes rapid, stimulus-specific structural and biochemical reorganization.

The mitochondria-to-cytosol volume percentage, representing volume occupancy, increased ∼60% from a basal 6.10 ± 1.24% to 9.85 ± 1.6% after just 1 minute of high glucose stimulation (Figure 1A, Table 1). Co-stimulation of Ex-4 alongside high glucose further amplified this effect, resulting in a larger increase of ∼90% to 11.53 ± 1.87% within 1 minute. These volume percentages began to decline across conditions by 5 minutes, which indicates temporal shifts consistent with mitochondrial swelling followed by fragmentation^56,57^. Cells stimulated with Ex-4 alone showed the most sustained and substantial expansion, with the mitochondria-to-cytosol volume percentage peaking at 15.06 ± 3.36% at 5 minutes. 3D renderings in Figure 1B illustrate a transition from the sparse, tubular network of unstimulated cells (grey) to the fragmented and swollen architecture induced by high glucose (red tones), while Ex-4 treatment, both alone and in combination with high glucose, promoted broader, increasingly integrated structures (green and blue tones). These illustrations offer insights into the structural changes occurring within the mitochondrial network.

**Figure 1:**
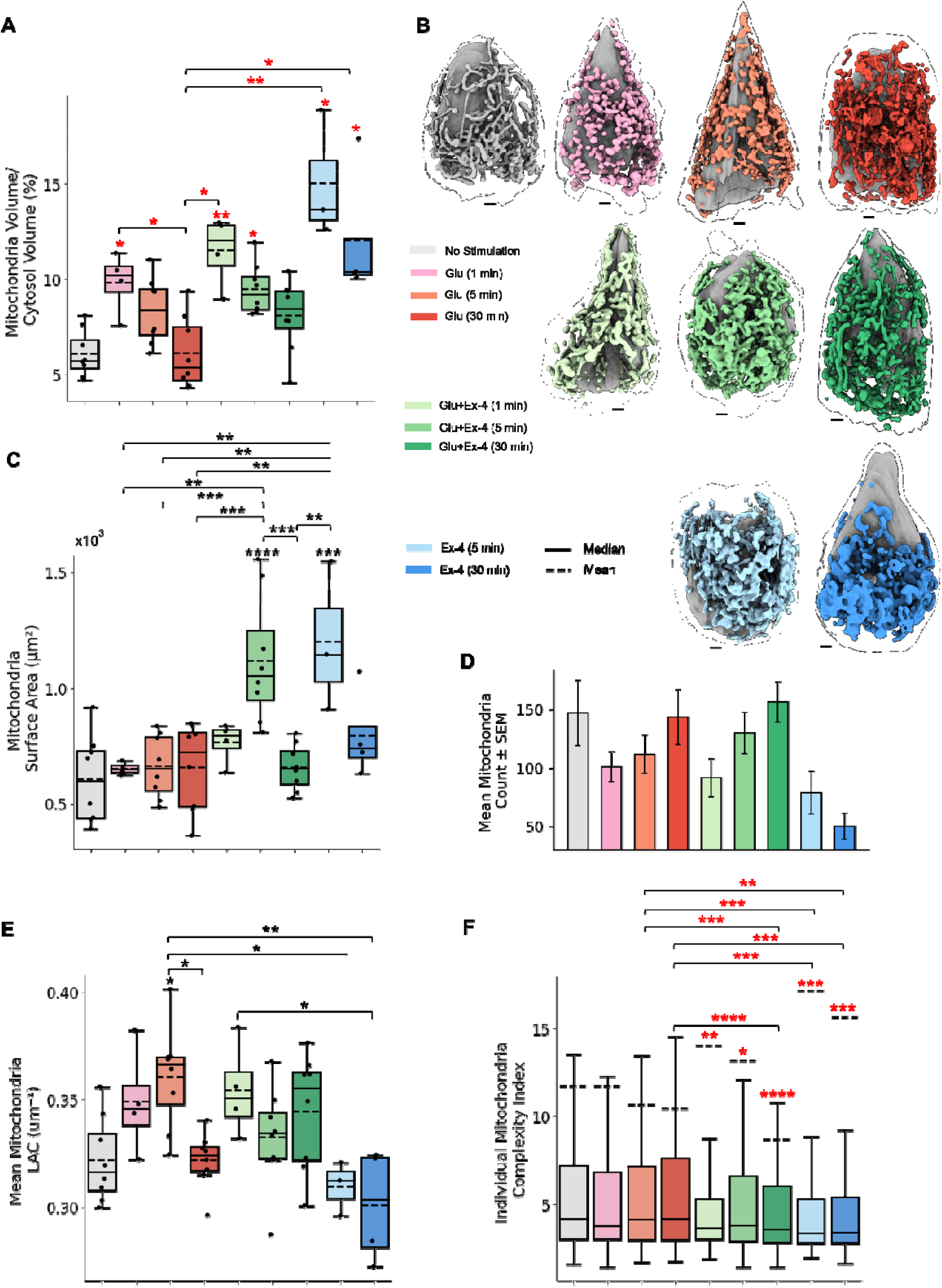
Quantitative assessment of whole-cell mitochondrial network remodeling. This figure illustrates the global structural and biochemical reorganization of the mitochondrial network across time and treatment conditions. (A) Mitochondria-to-cytosol volume percentage reveals organelle occupancy shifts, highlighting a rapid increase under early glucose and co-stimulation. (B) 3D illustrations generated using UCSF ChimeraX provide representative whole-cell reconstructions for each condition, visualizing the continuum from basal tubular networks to stimulus-induced fragmentation and swelling. (C) Mitochondrial surface area captures adaptive expansion, specifically identifying a significant rise unique to Ex-4 treatments at 5 minutes. (D) Individual mitochondrial count demonstrates a paradoxical decrease during early stimulation, suggesting initial organelle swelling rather than biogenesis. (E) Mitochondrial LAC (Linear Absorption Coefficient) maps voxel-level molecular density, showing rapid biochemical compaction under glucose stress. (F) Individual mitochondria complexity index quantifies structural intricacy, revealing that Ex-4 stabilizes higher network complexity compared to glucose alone. SEM stands for Standard Error of the Mean. Statistical significance relative to the unstimulated (NS) control is indicated by asterisks directly above individual condition whiskers, while statistical brackets denote significant pairwise differences between stimulated groups. Red asterisks represent Kruskal–Wallis tests with Dunn’s post-hoc comparison; black asterisks indicate one-way ANOVA followed by Tukey’s test (*p < 0.05, **p < 0.01, ***p < 0.001, ****p < 0.0001).

**Table 1:**
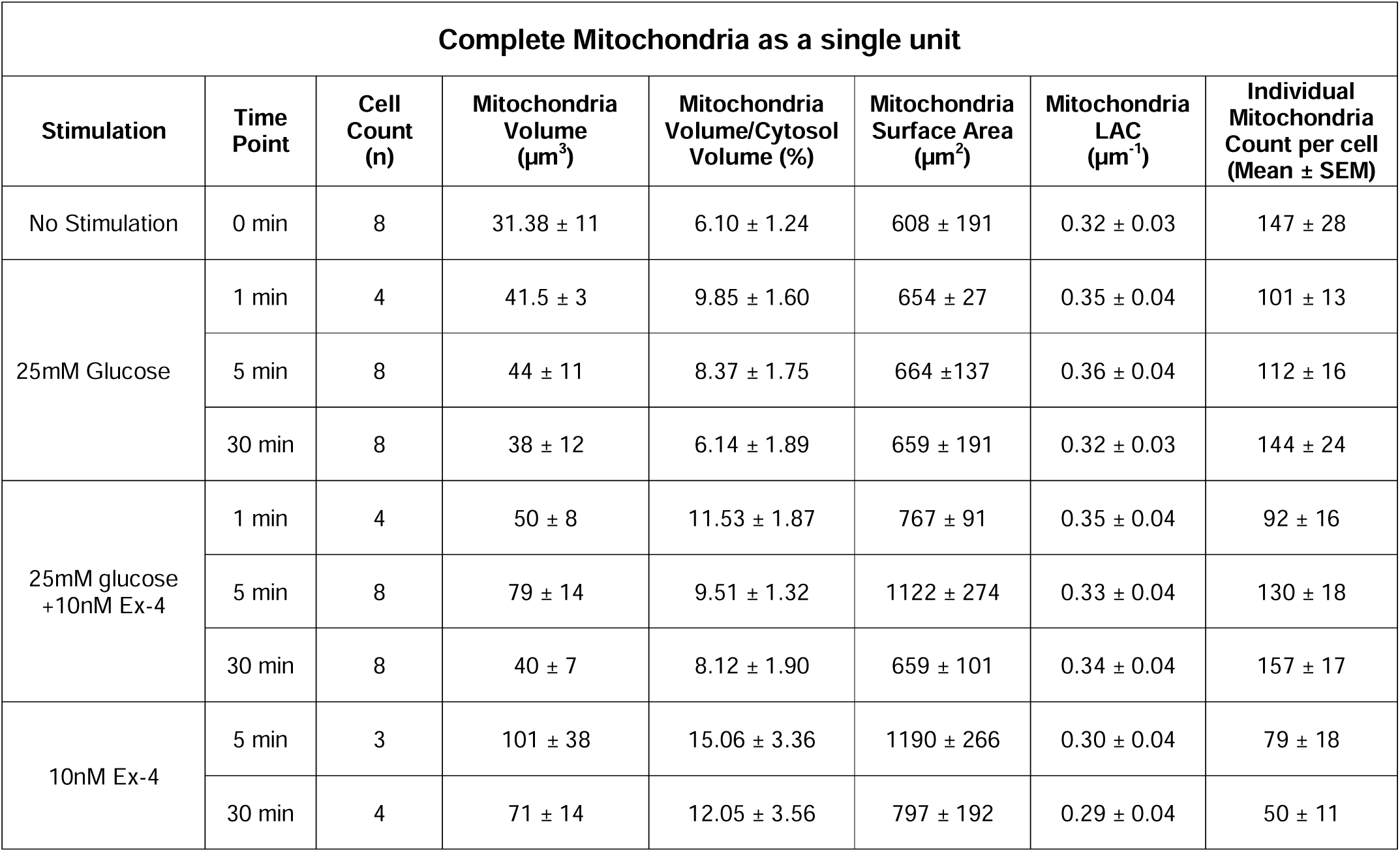
Quantitative metrics of global mitochondrial architecture. Summary of quantitative data from 55 whole-cell tomograms showing the statistical basis of the observed mitochondrial network remodeling. All metrics are reported as mean ± SD (standard deviation), except for individual mitochondria count, which is shown as Mean ± SEM (Standard Error of the Mean).

| Complete Mitochondria as a single unit |  |  |  |  |  |  |  |
| --- | --- | --- | --- | --- | --- | --- | --- |
| Stimulation | Time Point | Cell Count (n) | Mitochondria Volume ($\mu\text{m}^3$) | Mitochondria Volume/Cytosol Volume (%) | Mitochondria Surface Area ($\mu\text{m}^2$) | Mitochondria LAC ($\mu\text{m}^{-1}$) | Individual Mitochondria Count per cell (Mean ± SEM) |
| No Stimulation | 0 min | 8 | $31.38 \pm 11$ | $6.10 \pm 1.24$ | $608 \pm 191$ | $0.32 \pm 0.03$ | $147 \pm 28$ |
| 25mM Glucose | 1 min | 4 | $41.5 \pm 3$ | $9.85 \pm 1.60$ | $654 \pm 27$ | $0.35 \pm 0.04$ | $101 \pm 13$ |
| | 5 min | 8 | $44 \pm 11$ | $8.37 \pm 1.75$ | $664 \pm 137$ | $0.36 \pm 0.04$ | $112 \pm 16$ |
| | 30 min | 8 | $38 \pm 12$ | $6.14 \pm 1.89$ | $659 \pm 191$ | $0.32 \pm 0.03$ | $144 \pm 24$ |
| 25mM glucose +10nM Ex-4 | 1 min | 4 | $50 \pm 8$ | $11.53 \pm 1.87$ | $767 \pm 91$ | $0.35 \pm 0.04$ | $92 \pm 16$ |
| | 5 min | 8 | $79 \pm 14$ | $9.51 \pm 1.32$ | $1122 \pm 274$ | $0.33 \pm 0.04$ | $130 \pm 18$ |
| | 30 min | 8 | $40 \pm 7$ | $8.12 \pm 1.90$ | $659 \pm 101$ | $0.34 \pm 0.04$ | $157 \pm 17$ |
| 10nM Ex-4 | 5 min | 3 | $101 \pm 38$ | $15.06 \pm 3.36$ | $1190 \pm 266$ | $0.30 \pm 0.04$ | $79 \pm 18$ |
| | 30 min | 4 | $71 \pm 14$ | $12.05 \pm 3.56$ | $797 \pm 192$ | $0.29 \pm 0.04$ | $50 \pm 11$ |

Mitochondrial surface area analysis revealed differences that highlight the specific impact of Ex-4 (Figure 1C, Table 1). In the presence of Ex-4, significantly higher surface area values were observed at 5 minutes, both in the co-stimulation group (1122 ± 274 µm^2^) and with Ex-4 alone (1190 ± 266 µm^2^) compared to unstimulated cells (608 ± 191 µm^2^).

A notable structural paradox was identified when comparing mitochondrial volume occupancy to mitochondrial count. Although high glucose and co-stimulation with Ex-4 both led to a rapid increase in mitochondria-to-cytosol volume percentage at 1 and 5 minutes (Figure 1A), the mean count of individual mitochondria decreased during the same intervals compared to unstimulated cells (Figure 1D). As shown in Figure 1B, the early metabolic response is characterized by an “inflated” mitochondrial state, driven by organelle swelling in high glucose treated cells rather than the formation of new organelles. Also, for Ex-4 only treated cells the reduction in mean count of individual mitochondria can be attributed to increased fusion of the mitochondrial network^58–60^ (Figure 1B,D). In next sections, morphotype-specific analysis of mitochondria elucidates the subpopulation level structural remodeling in more detail.

SXT also provides quantitative estimates of biomolecular density through Linear Absorption Coefficient (LAC) values (voxel intensity value). Basal mitochondrial LAC in unstimulated cells (0.32 ± 0.03 µm^-1^) increased significantly following 5 minutes of high glucose stimulation (0.36 ± 0.04 µm^-1^), indicating rapid biochemical condensation immediately after stimulation (Figure 1E). In contrast, for cells treated with Ex-4 in absence of glucose, the average LAC value fall to 0.30 ± 0.04 µm^-1^ by 5 minutes and further to 0.29 ± 0.04 µm^-1^ by 30 minutes. We also analyzed the individual Mitochondrial Complexity Index to evaluate how these structural changes relate to network complexity. Higher complexity was observed in co-stimulated (Glu + Ex-4) cells at 1 and 5 minutes compared to glucose-only conditions, suggesting that Ex-4 supports maintenance of a more intricate mitochondrial network (Figure 1F).

### Morphotype-Specific Structural Responses

While global characterization provides a high-level overview of cellular state, accurate morphological phenotyping is essential to link structural states with metabolic function^1,61,62^. We therefore stratified the mitochondrial population into three distinct morphotypes based on individual volume: fragmented (< 0.06 µm^3^), intermediate (0.06 – 0.6 µm^3^) and interconnected (> 0.6 µm^3^) (Figure S1 and S2). Unlike the whole-cell metrics in the previous section, this analysis focuses on the structural footprint of a single mitochondrion within each morphotype. Our results reveal that mitochondria within these populations undergo different remodeling paths.

In fragmented population, under high-glucose stimulation, the average mitochondria-to-cytosol volume percentage increases significantly, reaching 0.006 ± 0.005 at 1 minute (Table 2, Figure 2A), while the mean number of fragmented mitochondria per cell decreases from a basal level of ∼102 to 48 (Table 2, Figure 2E). We also observed increase in mitochondrial surface area of fragmented populations by 1 and 5 minutes of high glucose stimulation (Figure S3). These changes appear to result from mitochondrial hypertrophy of fragmented population rather than an increase in mitochondrial number (Figure 2D). With prolonged glucose stimulation (30 minutes), this hypertrophic phase shifts toward increased fission, with the fragmented count rising to ∼77 per cell while the mean volume returns toward baseline (0.003 ± 0.003%).

**Figure 2:**
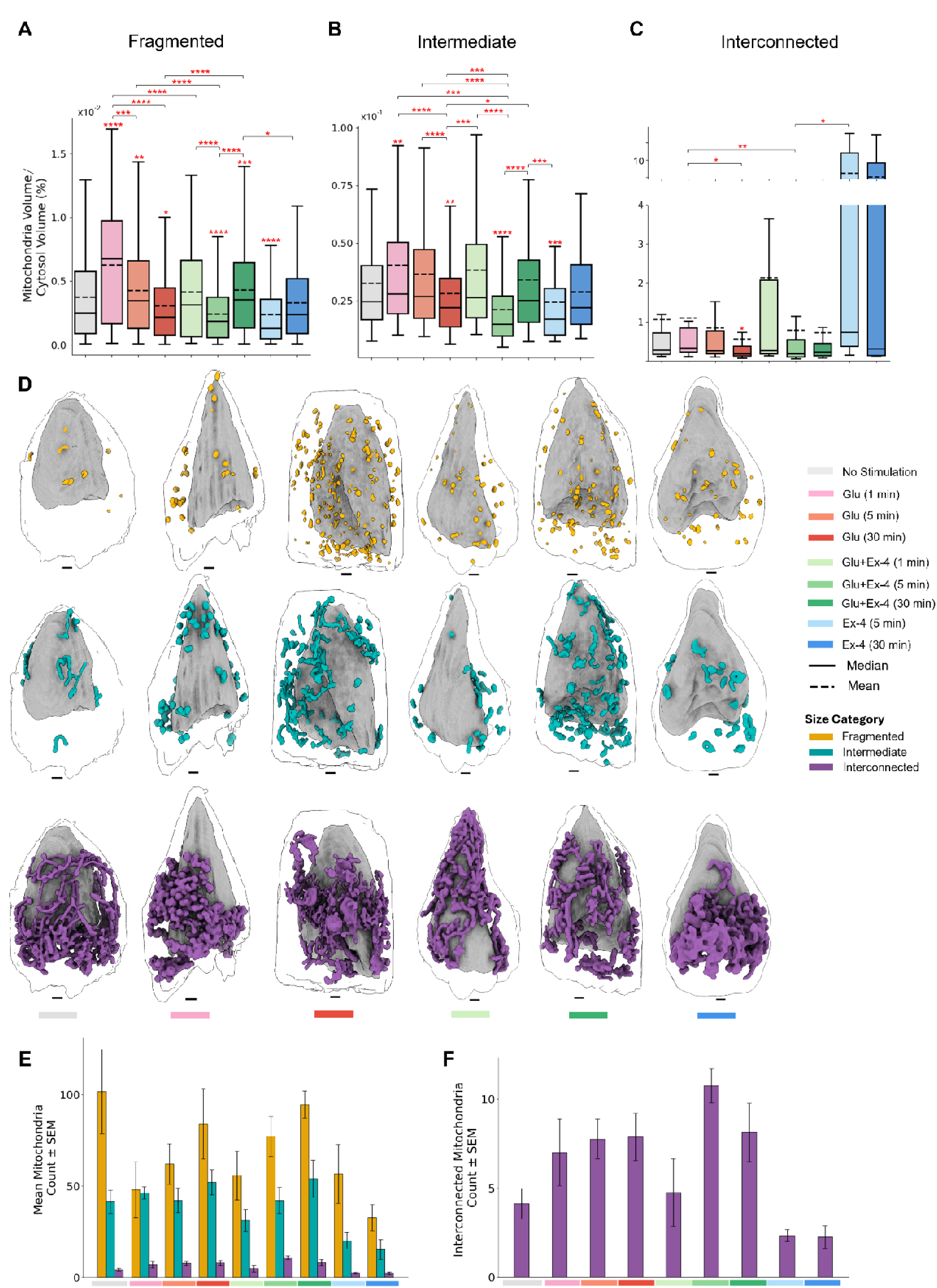
Morphotype-resolved analysis of mitochondrial populations. (A–C) Box plots show mitochondrial volume fractions normalized to cytosol for fragmented (A), intermediate (B), and interconnected (C) mitochondrial morphotypes. (D) Representative 3D whole-cell reconstructions illustrate spatial organization, color-coded by size category: fragmented (yellow), intermediate (cyan), and interconnected (purple). Illustrations were generated using UCSF ChimeraX. (E–F) Bar plots summarize the mean mitochondrial counts per cell for all morphotypes (E) and specifically for interconnected mitochondria (F). SEM stands for Standard Error of the Mean. Statistical significance relative to the unstimulated (NS) control is indicated by asterisks directly above individual condition bars/boxes, while horizontal lines denote significant pairwise differences between stimulated groups. Red asterisks indicate Kruskal–Wallis test with Dunn’s post hoc significance (*p < 0.05, **p < 0.01, ***p < 0.001, ****p < 0.0001).

**Table 2:**
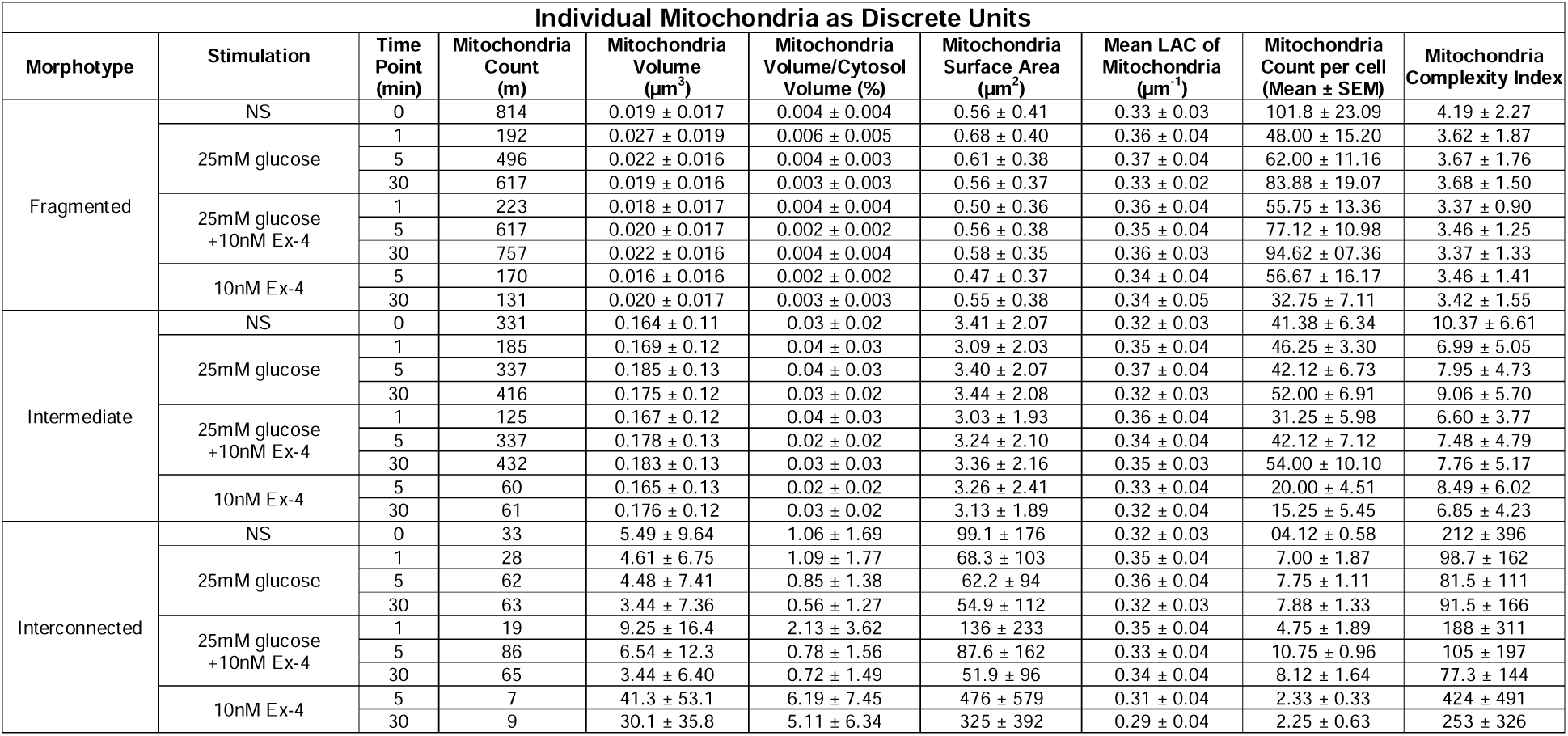
Morphotype-Specific Mitochondrial Metrics. Summary of the quantitative data for m = 6757 mitochondria across n = 55 whole-cell tomograms, providing the statistical basis for the morphotype-specific remodeling observed in the mitochondrial network. Each metric is mentioned as Mean ± S.D. (Standard Deviation), except for individual mitochondria count per cell which is quantified as Mean ± SEM (Standard Error of the Mean).

For intermediate and interconnected morphotype, a simultaneous increase is observed in both mean mitochondria-to-cytosol volume percentage and mean mitochondrial count at 1 min of high glucose stimulation (Figure 2B,C,E,F). All three morphotypes shows volume percentage increase at glucose 1 min, but the decrease in fragmented mitochondrial count is much more than the increase in intermediate and interconnected morphotype mean counts, resulting in “count paradox” observed in the previous section.

In contrast, co-stimulation with glucose and Ex-4 at 1 min, did not produce significant increases in individual volume percentage seen with glucose alone. Although the mean count of fragmented and intermediate mitochondria per cell still decreases, suggesting that Ex-4 limits glucose-induced swelling. This difference can also be seen in the 3D illustrations (Figure 2D), where fragmented (yellow), intermediate (cyan) and interconnected (purple) mitochondria appear less enlarged and more tubular than those treated with glucose alone. This structural stability is consistent with the known protective effects of incretin signaling on mitochondrial dynamics^58,63,64^.

While fragmented and intermediate mitochondria showed volume stabilization with Ex-4, a different behavior was observed in the interconnected population. During prolonged high glucose exposure (30 min), the mean volume fraction of interconnected mitochondria significantly decreases to 0.56 ± 1.27% from a basal level of 1.06 ± 1.66% (Figure 2C, Table 2), suggesting partial collapse of the fused network. In contrast, Ex-4 alone results in a massive expansion in mean volume occupancy, reaching 6.19 ± 7.45% at 5 min and 5.11 ± 6.34% at 30 min (Table 2 and Figure 2C), even though approx. two interconnected mitochondria were present per cell. This confirms that Ex-4 promotes a hyper-fused state that consolidates the dispersed mitochondrial population into a massive, high-complexity network. Interestingly, the Ex-4 in co-stimulation partially preserves the collapse of interconnected mitochondria induced by high glucose by 30 minutes (0.72 ± 1.49%, Table 2).

Overall, we observe that a small number of large and complex structures of interconnected morphotype can disproportionately influence total mitochondria analysis, often masking the underlying morphotype-specific patterns.

### Unsupervised Geometric Fingerprinting of Intermediate Morphologies

While in previous section we identified significant remodeling across all morphotypes, the intermediate morphotype represents a critical transitional state between fragmented and interconnected. As shown in illustrations in Figure 2D, fragmented mitochondria (< 0.06 µm^3^) are generally too small and structurally uniform for deeper geometric classification, while interconnected mitochondria (> 0.6 µm^3^) are often too large and topologically complex to resolve into distinct sub-types. Consequently, we focused this section specifically on the intermediate population to explore the diversity of this “morphological buffer”. By using quantitative shape-descriptive features, such as, volume, tortuosity, sphericity and flatness (Table S1), we projected each intermediate mitochondria into 2-dimensional space, identifying three distinct sub-clusters using unsupervised k-means clustering (Figure 3A). This reveals that the intermediate population is a continuum of three geometric sub-clusters.

**Figure 3:**
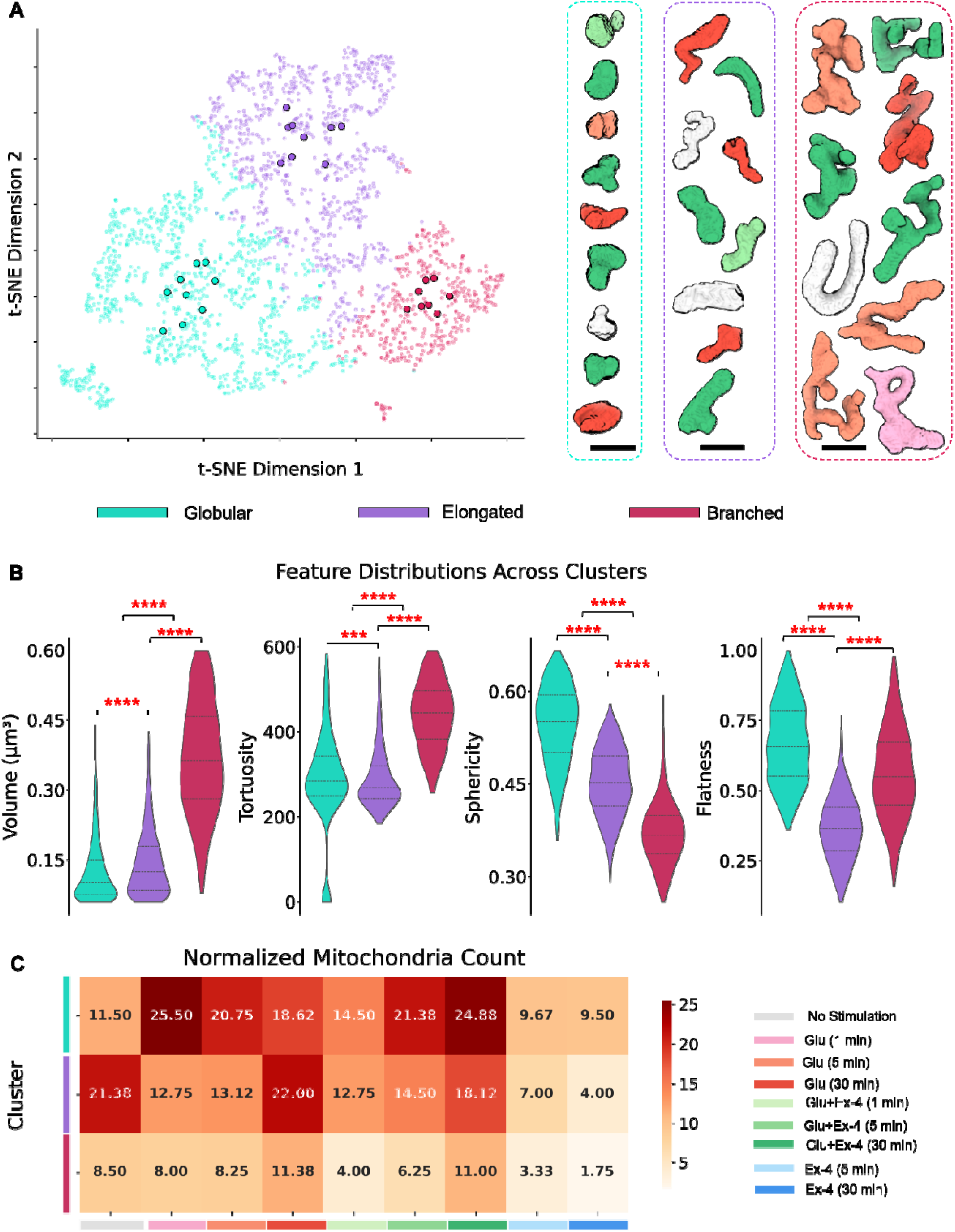
Unsupervised classification of intermediate mitochondria. (A) A 2D t-SNE visualization of the intermediate mitochondrial population (0.06 – 0.6 µm^3^), where individual mitochondria are clustered via k-means based on multi-dimensional shape features. Highlighted data points correspond to representative 3D renderings on the right, which are color-coded by cluster identity: Globular (cyan), Elongated (purple), and Branched (red). Mitochondria are colored based on treatment conditions, as shown in legend (C). (B) Violin plots illustrating the distribution of key geometric parameters, Volume, Tortuosity, Sphericity, and Flatness, across the three clusters. These distributions define the distinct structural “fingerprints” of the intermediate pool, ranging from compact, high-sphericity globular units to high-tortuosity branched networks. Red asterisks indicate Kruskal–Wallis test with Dunn’s post hoc significance (*p < 0.05, **p < 0.01, ***p < 0.001, ****p < 0.0001). (C) Normalized mitochondrial counts per cell for each cluster across stimulation conditions and time points. The color intensity reflects the abundance of specific morphotypes, highlighting the rapid globular shift under acute glucose stress and the stabilizing effect of Ex-4, which delays fragmentation and promotes consolidated network architecture.

The first sub-cluster, Globular-intermediate, is characterized by the highest sphericity and flatness (Figure 3B). These represent the “hypertrophic” units, like fragmented morphotype, that swell during early glucose stress. The second is an Elongated-intermediate cluster, which represents the baseline tubular state of the mitochondrial network. Lastly, we observed a Branched-intermediate sub-cluster with highest volume and tortuosity (Figure 3B). This cluster potentially serves as the structural building blocks for larger interconnected networks. Example geometries from the three sub-clusters are illustrated in Figure 3A.

The normalized counts of these sub-clusters reveal how glucose and Ex-4 “rewire” the network’s transitional pool (Figure 3C). Under high glucose alone, the Globular-intermediate mean count increases rapidly within 1 minute, rising from 11.50/cell at baseline to 25.50/cell. This change likely reflects an acute stress response, where intermediate mitochondria shift toward a more rounded and swollen morphology as observed in previous section (Figure 2). After 30 minutes of glucose exposure, the Elongated-intermediate dominates the distribution again indicating an adaptive remodeling response.

Interestingly, with co-stimulation the rapid rise in the Globular-intermediate population is much less pronounced (14.50 per cell). The co-stimulation delays the abundance peak of Globular-intermediate, reaching levels comparable to 1 min glucose spike after 30 minutes of sustained exposure. This suggests that Ex-4 acts as a structural stabilizer that prevents the “shock” of high glucose from immediately forcing the intermediate population into a swollen, globular state.

### Morphotype-Specific Spatial Partitioning and Radial Enrichment

To determine whether structural remodeling is coupled with changes in organelle localization, we performed a contour-based radial enrichment analysis (observed-to-expected distribution) along a normalized axis from the nuclear membrane to the cell periphery binned into spatial contours (Figure 4A). Differences in mitochondrial spatial organization between stimulation conditions were quantified using the Wasserstein distance between enrichment distributions across contour bins (Figure S4). Like previous sections, we performed morphotype-specific enrichment analysis to prevent losing spatial patterns of fragmented and intermediate populations, as the interconnected morphotype dominates the total mitochondrial volume. Generally, morphotypes display a central-cytoplasmic enrichment towards high microtubule density and metabolic demand^65,66^, with significant depletion near the nucleus and the cell membrane (Figure 4B).

**Figure 4:**
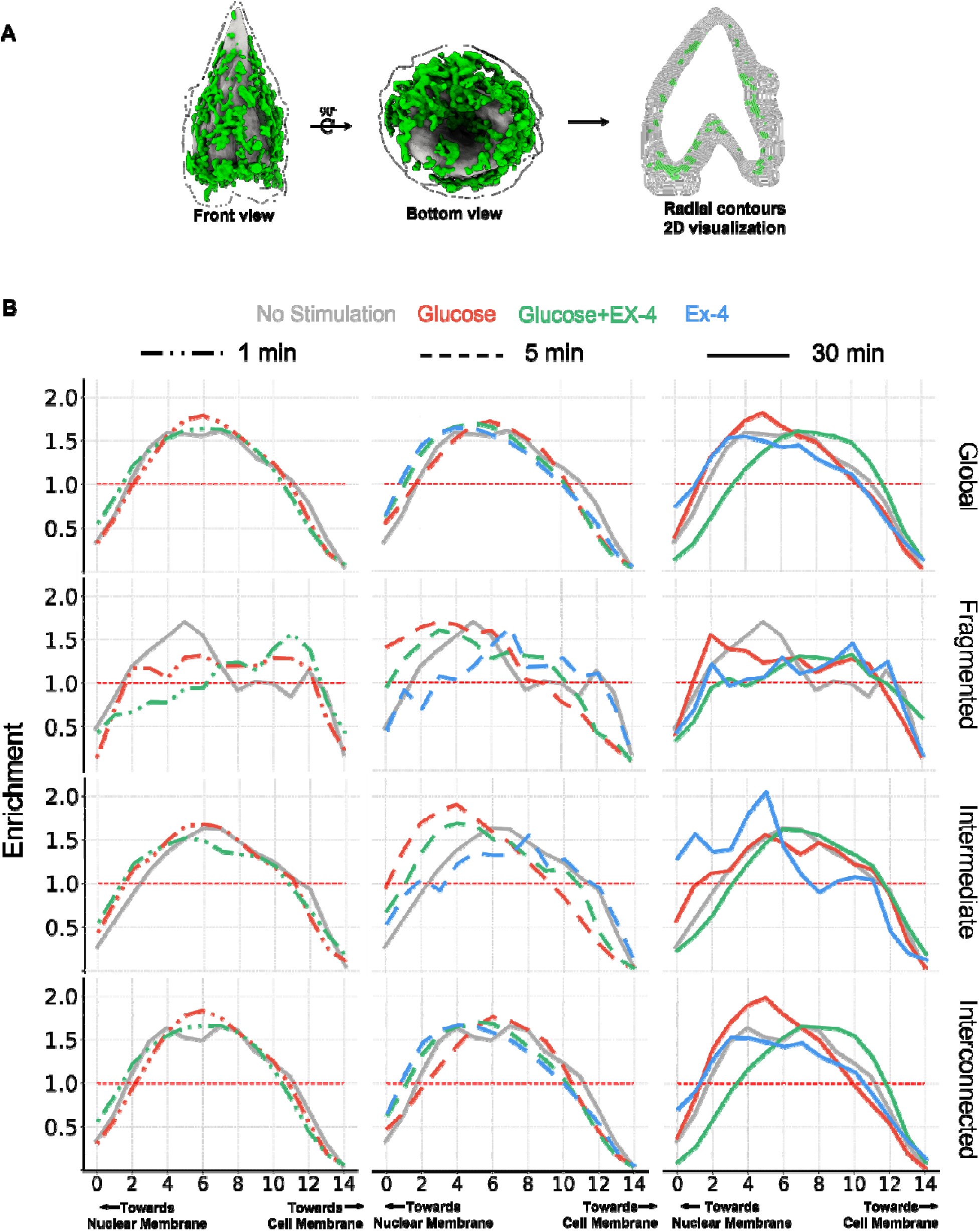
Contour-based Radial Profiling and Spatial Enrichment of Mitochondrial Morphotypes. (A) Front and bottom views of representative 3D whole-cell reconstructions visualized using UCSF ChimeraX, along with a 2D slice illustrating the radial contours. The cytoplasm is partitioned into spatial contours starting from the nuclear membrane (contour 0) to the cell membrane (contour 14) (B) Mitochondrial enrichment (Observed-to-Expected ratio) across spatial contours. Rows correspond to the Global (Complete Network), Fragmented, Intermediate and Interconnected populations, while columns indicate time points: 1 min (left, dash-dot), 5 min (middle, dash) and 30 min (right, solid). The horizontal red short-dash line at y=1.0 depicts no observed enrichment or depletion as compared to expected.

At 1 min, fragmented mitochondria show high spatial reorganization. In high glucose-treated cells, fragmented morphotypes lose the peak (contours 4-6, gray line) and redistribute uniformly across the cytoplasm (Figure 4B: left column, second row, red dash-dot line). Co-stimulation with Exendin-4 further shifts the distribution towards outer cytoplasmic and peripheral contours (10-12, green dash-dot line). Such fragmentation is a well-established stress phenotype that produces motile fragmented units responding to Ca^2+^ signaling or metabolic fluctuations^1,67^. In contrast, the intermediate and interconnected morphotype shows no major spatial shifts at 1 min time point (Figure 4B: left column, third row).

By 5 min, both fragmented and intermediate morphotypes show increased localized perinuclear enrichment under high glucose alone and co-stimulation. This likely stems from fission of interconnected mitochondria to fragmented and intermediate morphotypes, as interconnected volume simultaneously decreases in volume fraction (Table 2, Figure 2C). Notably, as Ex-4 antagonizes glucose induced fission, the perinuclear enrichment under co-stimulation is significantly lower compared to glucose alone (Figure 4B, middle column, second and third row, contours 0-6). This can be further validated by the Ex-4 only treatment, which shows no perinuclear enrichment either of fragmented or intermediate morphotypes in the absence of hyperglycemic stress.

By 30 min, all morphotype profiles under glucose only partially return toward the no-stimulation state, suggesting a recovery from early fission rather than persistent fragmentation. Conversely, co-stimulation promotes sustained spatial shift of intermediate and more so of interconnected morphotypes toward the cell periphery (Figure 2B, right column, fourth row). Overall, these results indicate that mitochondrial positioning is dynamically regulated by both stimulation time and mitochondrial morphotype, where early glucose exposure triggers localized perinuclear fission, while Ex-4 signaling preserves network structure and shifts mitochondrial distribution towards cell periphery.

### Spatiotemporal Mapping of Mitochondrial Metabolic Density

To examine structural remodeling association with shifts in metabolic output, we mapped mitochondrial Linear Absorption Coefficient (LAC) values across the nuclear–to–cell membrane axis, as done for enrichment (Figure 4A). As LAC scales with biomolecular density, it can be used as an indicator of macromolecular packing and the metabolic state of the organelle^46,48,51,52,68^. In mitochondria, increased LAC typically signifies higher metabolic density and ATP production potential, as the matrix is the primary site for the TCA cycle and oxidative phosphorylation.

At 1-minute, co-stimulated (Glu + Ex-4) cells already show elevated LAC values across all morphotypes, particularly toward the cell periphery (contours 10–14) (Figure 5, left column, green). In contrast, glucose-only treatment reaches comparable maximal LAC levels later at 5 minutes (Figure 5, middle column, red), as also illustrated by the representative 3D renderings in Figure 6. This suggests that Ex-4 accelerates the overall metabolic shift, aligning with the studies that incretins like GLP-1/Ex-4 enable β-cells to become active more rapidly^63,69,70^.

**Figure 5:**
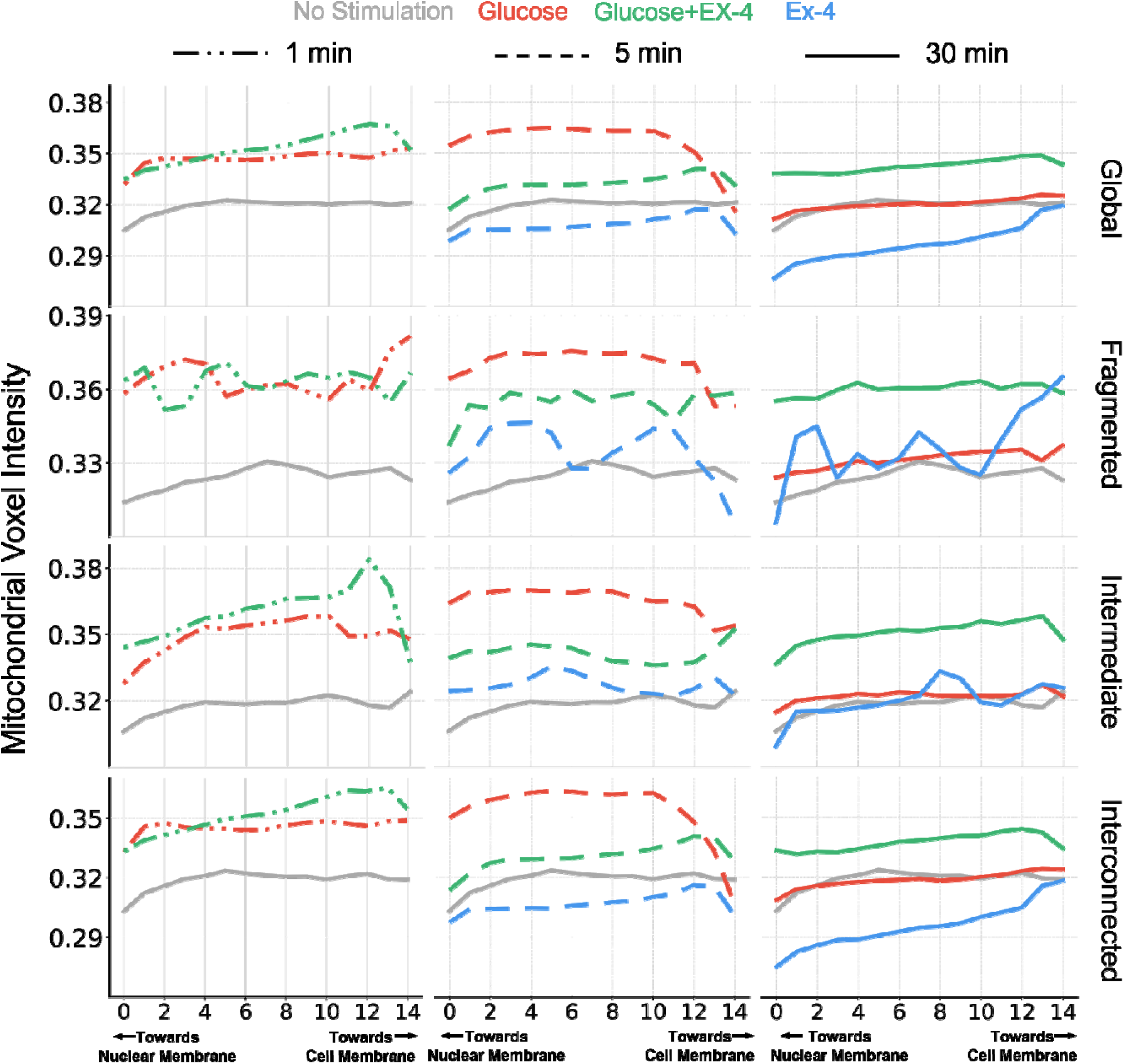
Mitochondrial voxel-level metabolic intensity (LAC intensity) across sub-cellular spatial bins for four major morphological states. A) Spatial distribution of mitochondrial LAC value across the cytoplasmic contours. Columns represent temporal progression (left: 1 min; middle: 5 min; right: 30 min), and rows correspond to mitochondrial morphological classes. Within each subplot, color denotes stimulation condition: No Stimulation (grey); Glucose (red); Glucose + Ex-4 (green); and Ex-4 (blue). Line styles indicate time points: 1 min (dash-dotted); 5 min (dashed); and 30 min (solid). The x-axis represents subcellular radial bins extending from the nuclear membrane toward the cell membrane, and the y-axis shows the mean mitochondrial voxel intensity within each radial bin.

**Figure 6:**
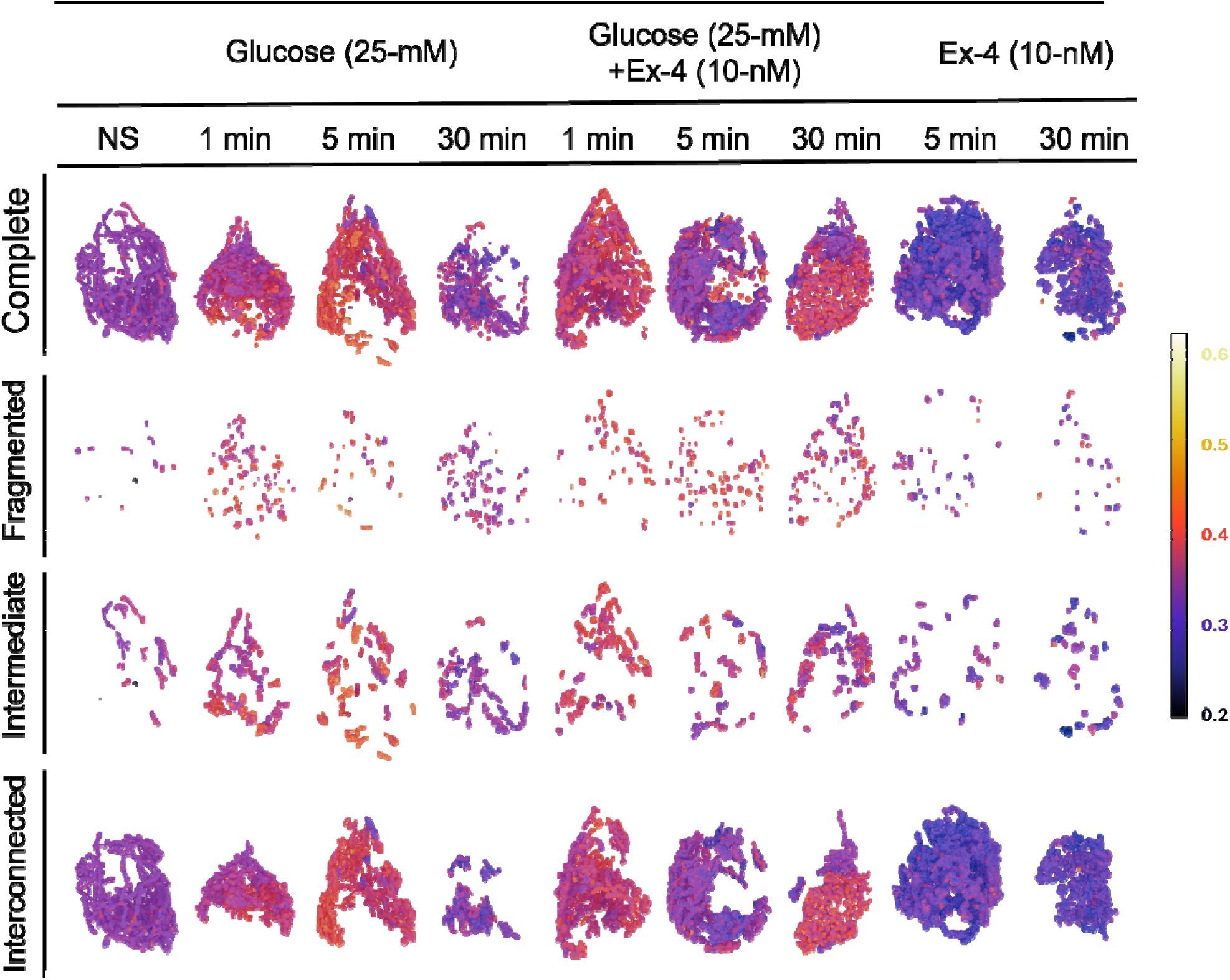
Stimulus- and morphology-dependent modulation of mitochondria. Representative mitochondrial LAC maps categorized by morphotype and stimulation condition. Rows indicate morphotypes and columns indicate treatment conditions and time points. Color scale shows low (dark) to high (warm) LAC values, highlighting stimulus-dependent mitochondrial remodeling.

Interestingly, intermediate and interconnected morphotypes in co-stimulated cells show relatively high LAC values near cell membrane regions compared to perinuclear regions at 1 and 30 minutes (Figure 5, left and right column). This observation is notable because Ex-4 also promotes mitochondrial redistribution toward the cell periphery. The presence of metabolically dense mitochondria in this region may help meet the local energy demand required for insulin secretion. By 30 minutes, only the co-stimulated condition maintained higher LAC values across morphotypes compared with control and glucose-only conditions (Figure 5, right column, green line and Figure S5, last row). This prolonged activity is visualized in Figure 6, where co-stimulated mitochondria retain a distinct “red shift” in intensity (higher LAC), while glucose-only cells transition toward lower density tones.

Ex-4 treatment alone displays a markedly different pattern. While fragmented and intermediate morphotypes show a minor LAC increase at 5 minutes, interconnected mitochondria exhibit a progressive decrease in metabolic density that deviates further from the no stimulation by 30 minutes (Figure 5 last row, Figure S5 last row and Figure 6 “blue shift” in last two columns). This confirms that the densification effect of Ex-4 is glucose-dependent and without glucose availability the metabolic density falls, pointing to the link between LAC and metabolic activity.

Importantly, our spatial binning strategy reveals microenvironment-specific metabolic signatures that would be obscured by traditional global intensity metrics or 2D projections.

## DISCUSSION

### The Structural Continuum

The above reported results suggest that classifying mitochondria simply as “fused” or “fragmented” does not fully capture the structural changes occurring within the cell. Mitochondrial stratification into fragmented, intermediate, and interconnected groups showed an early hypertrophic phase within 1 minute of high-glucose treatment. This is reflected in the observed “count paradox”, where total mitochondrial volume increases despite a reduction in organelle number. This swelling likely represents a state of metabolic overload and transient membrane depolarization^66,71–74^, preceding the high-frequency fission events observed at the 30-minute mark.

The unsupervised geometric fingerprinting of the intermediate morphotype further reveals that this population serves as a “morphological buffer.” The shift from elongated to globular-intermediate sub-clusters under glucose stress reflects alterations in fission–fusion balance, stress adaptation, and bioenergetic demand^1,65,75^.

### Morphotype-Resolved Bioenergetics

The analysis of metabolic signatures at individual morphotype level using Linear Absorption Coefficient (LAC) allowed to quantify biomolecular density, moving beyond the shape-based analysis. The whole-network LAC intensity profiles appeared stable because interconnected mitochondria dominate the total voxel pool, as their high metabolic intensity and superior ATP-generating capacity^71–73^ masks the dysfunction occurring in fragmented and intermediate mitochondria. The network deconstruction revealed a distinct morphotype-specific metabolic pattern. Fragmented mitochondria showed significantly reduced and variable LAC intensities, particularly near the nucleus under glucose stress, consistent with early dysfunction and reduced oxidative phosphorylation^66,74^. In contrast, interconnected mitochondria maintained consistently higher LAC values across radial contours, supporting their relatively stable metabolic state^72,73^. This type of analysis helps address some of the limitations of earlier fluorescence-based and 2D imaging approaches^76^.

### Exendin-4 as a Structural and Bioenergetic Stabilizer

One of the central observations emerging from this study is the multi-dimensional stabilizing effect exerted by Exendin-4 (Ex-4). A hyper-fused mitochondrial configuration instead becomes prominent when Ex-4 is present, consolidating mitochondria into massive and highly complex interconnected networks, whereas high glucose alone drives the system toward fragmentation and structural collapse. This preservation of mitochondrial structure is consistent with the well-established protective roles of GLP-1 receptor agonists in limiting glucotoxicity^63,64^.

Increased LAC values, reflected as a noticeable “red shift” in the co-stimulated cells, further suggest that Ex-4 does not merely maintain mitochondrial size but also preserves their biochemical density. Within mitochondria, LAC generally scales with the degree of macromolecular packing inside the matrix, particularly involving enzymes of the TCA cycle together with components of the electron transport chain. A higher metabolic potential, therefore, becomes apparent from the Ex-4-induced increase in LAC, indicating strengthened oxidative phosphorylation capacity and effectively priming the β-cell to sustain prolonged insulin demand under continued metabolic stimulation.

### Structure–Density–Location Coupling

The spatial distribution of mitochondrial morphotypes together with their LAC values adds a new dimension to the understanding of β-cell polarity. We observed that Ex-4 promotes a localized increase in metabolic density specifically near the cell periphery. Near the plasma membrane, the positioning of dense interconnected mitochondria creates localized ATP-generating hubs that help support the high energy demand of insulin vesicle docking and exocytosis. Similar spatial polarity has also been described in the healthy β-cells, where mitochondria located near the cell membrane show dense cristae structures^77–79^, showing the protective effect of Ex-4.

In contrast, fragmented and low-density mitochondria tend to move towards the nuclear periphery when cells are stimulated with glucose alone. This accumulation of fragmented mitochondria near the nucleus likely reflects a stress response in which dysfunctional organelles are isolated for mitophagy^80,81^.

Taken together, these divergent spatial signatures point to a structure-density-location coupling in the mitochondrial network, i.e., morphotype, macromolecular packing and radial position do not vary independently but shift in concert during metabolic remodeling. Under glucose stress, fragmented morphotypes with low LAC accumulate perinuclearly, whereas under Ex-4 co-stimulation, interconnected morphotypes with high LAC are enriched at the cell periphery. This coupling suggests that the metabolic state of a mitochondrion is jointly encoded by its shape, its biochemical density, and where it resides within the cell, rather than by any one parameter in isolation.

### Open Questions and Future Directions

Our findings open new avenues for structure-based therapeutic design. Beyond simply inhibiting fission, future interventions could be engineered to “tune” the mitochondrial morphology, density and location, targeting matrix condensation or cristae stabilization to restore the high-density, interconnected morphotypes observed with Ex-4.

Future research must determine if Ex-4-induced “morphological memory” persists under chronic stress and identify the molecular triggers linking GLP-1 signaling to matrix compaction. Understanding these mechanisms will be essential for translating 3D morphometric insights into targeted clinical treatments. More broadly, whether the structure–density–location coupling observed here extends to other organelle systems or cell types remains an open question. This high-fidelity dataset and the comprehensive analysis of mitochondria also offer organelle-level spatiotemporal and density distributions for whole-cell modeling.

## Resource availability

### Lead Contact

Requests for further information and resources should be directed to and will be fulfilled by the lead contact, Jitin Singla.

### Materials availability

This study did not generate new unique reagents.

### Data and Code Availability

The scripts used for this study are available at: https://github.com/jitinsingla/MitoMorphologySXT. Additional information required to reproduce or extend this work can also be obtained from the corresponding author.

## Acknowledgments

This work was supported by funding from the Chellaram Diabetes Research Centre (CDRC). We also acknowledge IIT Roorkee for financial support and access to computational facilities. We thank Valentina Loconte for assistance with data collection.

## Author Contributions

J.S. conceived and designed the study. K.L.W. performed the cell culture stimulation experiments and carried out data segmentation. A.Y., A.S., A.D., and R.V. performed the formal data analysis. A.S. and A.S. also contributed to developing the analysis pipeline and assisted with molecular density visualization using Python. A.S. prepared the initial draft of the manuscript, and all authors contributed to revisions. A.D., J.S., and K.L.W. reviewed and edited the final manuscript.

## Declaration of generative AI and AI-assisted technologies

During the preparation of this work, the author(s) used Gemini3 for editing the text to modify the sentence flow. In Graphical abstract, mitochondrial network illustration was also generated using Gemini3. After using this tool or service, the author(s) reviewed and edited the content as needed and take(s) full responsibility for the content of the publication.

## Supplementary Information

Document S1. Table S1 and Figures S1–S6

## STAR METHODS

### Deposited Data

| RESOURCE | SOURCE | IDENTIFIER |
| --- | --- | --- |
| Raw and analyzed datasets from unstimulated cells, cells stimulated with 25 mM glucose for 5 and 30 minutes, and cells co-stimulated with glucose and Ex-4. | White et al., 2020<sup>52</sup> | N/A |
| Raw and processed data from cells stimulated with 25 mM glucose (1 min), co-stimulated with glucose and Ex-4, and treated with 10 nM Ex-4 for 5 and 30 min. | Loconte et al., 2022<sup>48</sup> | N/A |

Experimental Models: Cell Lines
| RESOURCE | SOURCE | IDENTIFIER |
| --- | --- | --- |
| INS-1E | Addex Bio | Cat# C0018009; RRID: CVCL_0351 |

Software and Algorithms
| RESOURCE | SOURCE | IDENTIFIER |
| --- | --- | --- |
| ImageJ-Fiji 1.54f | Schindelin et al. (2012)<sup>82</sup> | https://imagej.net/ij/ |
| IMOD 4.10.42 | Kremer et al. (1996)<sup>83</sup> | https://bio3d.colorado.edu/imod/ |
| ChimeraX | Elaine C. Meng et al. (2023)<sup>84</sup> |  |
| Organelle identification pipeline | White et al. (2020)<sup>52</sup> | N/A |
| Individual organelle identification pipeline | This paper | N/A |
| Morphometry | This paper | N/A |
| Spatial localization of molecular density (LAC) | This paper | N/A |
| Spatiotemporal localization of mitochondria using contours | This paper | N/A |

### Methods

#### Soft X-ray Tomography

Soft X-ray tomography (SXT) data were acquired at 517 eV using the XM-2 soft X-ray microscope at the National Center for X-ray Tomography, Advanced Light Source (Lawrence Berkeley National Laboratory), as previously described by Le Gros et al. (2014)^85^. Cells were imaged using a zone plate objective and maintained under cryogenic helium conditions to preserve native ultrastructure. Projection images were collected over a full 180° rotation in 2° increments. Tomograms were reconstructed from the acquired projections using the AREC3D^86^ iterative reconstruction method, generating three-dimensional datasets with near-isotropic spatial resolution determined by the 60-nm zone plate objective. Voxel intensities were normalized to obtain quantitative linear absorption coefficient (LAC) values across all samples.

#### Dataset

Soft X-ray tomograms of Rat insulinoma INS-1E cells were obtained from previously published datasets Loconte et al. (2022)^48^ and White et al. (2020)^52^. The dataset included reconstructed tomograms and corresponding manually segmented masks. Following the quality assessment, 55 out of 56 cells were used in this study (Figure S2).

#### Identification of individual organelles

To quantify properties at the single-organelle level, connected-component labeling was applied to the binary whole-cell organelle mask using 26-connectivity in three-dimensional space, implemented using the *scipy.ndimage.label* function. Under this connectivity criterion, voxels sharing faces, edges, or corners were considered part of the same organelle. The number of individual organelles was defined as the total number of connected components detected in the labeled mask. To ensure methodological reliability detection of individual organelles from few whole-cell masks were manually verified using ImageJ^82^.

#### Absolute Volume and Surface Area calculation

For each mitochondrial mask (either complete whole-cell mask or individual mitochondrial mask), absolute volume in µm^3^ was computed as *V* = *N_voxels_* x *l*^3^, where *l* is the voxel edge length in micrometers. The voxel edge length is computed from imaging parameter called lac factor which is specific to each cell.

Surface area was estimated directly using a face-exposure method. For each voxel belonging to the mitochondria, all six orthogonal faces (±x, ±y, ±z) were examined. If a voxel face does not make interface with the same mitochondria, it is counted towards surface area. Finally absolute surface area in µm^2^ was computed as *SA* = *N_faces_* x *l*^2^.

#### Mitochondrial Complexity Index

Mitochondrial morphology ranges from compact, spherical structures to highly elongated, branched, and interconnected networks. While volume reflects mitochondrial mass, surface area captures membrane expansion associated with elongation and branching. The MCI integrates volume and surface area, providing a scale-independent measure of morphological complexity. Small, compact spherical organelles have low MCI compared to organelles with complex topology.

For each individual mitochondrion, the MCI was calculated as:

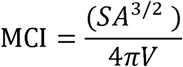

where, *SA* is the mitochondrial surface area (µm²) and *V* is the mitochondrial volume (µm³)^87,88^.

#### Clustering-based identification of mitochondrial morphologies

A total of 8 shape-descriptive features (Table S1) were calculated from 2284 intermediate size mitochondria. Using k-means clustering on the shape features, intermediate mitochondria were segregated into 3 clusters. The distribution of features and corresponding clusters were visualized using t-SNE embeddings. Based on the features and visual inspection the three clusters were named Globular, Elongated and Branched.

#### Radial Contour Mapping

To quantify the intracellular spatial organization of mitochondria, a normalized radial coordinate system was established within the cytoplasmic compartment. For each cytoplasmic voxel, Euclidean distance transforms were computed using SciPy to determine the shortest distance from the nuclear (*d_nuc_*) and cell boundaries (*d_cell_*). A normalized radial position was then calculated as 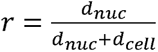. As *r* values rise from 0 to 1, the voxel location changes from nuclear to cell boundary. This normalization converts intracellular space into a morphology-independent radial axis, allowing comparison between cells with different sizes and shapes.

The cytoplasm was subsequently subdivided into 15 concentric radial contours by discretizing the normalized radial coordinate into equal intervals. The contour *c*∈[0, 14] contains the cytoplasmic region within range 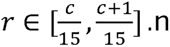 Each contour therefore represents a fractional spatial shell extending progressively from the perinuclear region toward the cell periphery.

#### Enrichment Analysis

For individual cell, the observed mitochondrial distribution was determined by quantifying mitochondrial voxels located within each contour as 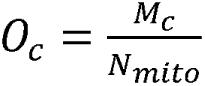, where *M_c_* corresponds to mitochondrial voxel counts within the contour *c*, and *N_mito_* epresents the total number of mitochondrial voxels in the cell. Because cytoplasmic volume is not uniformly distributed across contours, the observed distribution has a volumetric bias, i.e. contours towards cell membrane will contain more mitochondrial voxels due to more available cytoplasmic space. Therefore, we normalized the observed distribution with the expected distribution based on cytoplasmic volumes. For each contour, the expected distribution was computed as 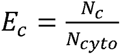, where *N_c_* represents the number of cytoplasmic voxels within the contour *c*, and *N_cyto_* denotes the total cytoplasmic voxel count in the cell. *E_c_* reflects the probability of mitochondrial localization assuming random spatial occupancy.

Finally mitochondrial enrichment, i.e., observed-to-expected ratio for each contour was computed as 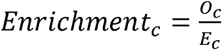. An enrichment value greater than one indicates preferential mitochondrial accumulation within a given radial region, whereas values below one indicates spatial depletion relative to cytoplasmic volume availability. This normalization enables quantitative comparison of mitochondrial positioning independent of cellular morphology or cytoplasmic volume heterogeneity. Eventually, to compute enrichment for a single treatment or condition, the enrichment was averaged across cells for each contour.

#### LAC-based Spatial Density Analysis

To investigate spatial variations in mitochondrial intensity, voxel-level LAC measurements were extracted directly from reconstructed tomograms. The LACs were then mapped to cytoplasmic contours based on the normalized radial position of the voxel. Then the spatial profile of LAC-based density was computed by averaging all the voxel LAC values within each contour.

#### LAC-based 3D Colormaps

The mitochondrial LAC values across all the cells combined were normalized between 0.2 and 0.6 to ensure equal color mapping across drug and time conditions. The Voxel-Gen Tools Blender^89^ add-on is used to generate 3D structures^90^. It takes the binary and LAC-normalized arrays as input and generates a voxel-mesh, where each cube is colored according to the original image values. Smoothing is achieved through volumetric triangulation using Geometry Nodes and scripting, with LAC values assigned to individual vertices based on the minimum Euclidean distance transform (EDT). 3D environment lighting and camera effects are implemented to generate the final render. The CMRMap from the Matplotlib library is used for final plot.

#### Graph-based Network Analysis

To characterize the topological organization of individual mitochondria, each mitochondrial mask was reduced to its medial axis by three-dimensional skeletonization using the skimage.morphology.skeletonize_3d function. For every skeleton voxel, the number of 26-connected neighbours was determined by convolution with a 3×3×3 kernel (scipy.ndimage.convolve). End points were defined as skeleton voxels with a single neighbour and branch points as voxels with three or more neighbours, the latter grouped into discrete junctions through connected-component labeling (scipy.ndimage.label, 26-connectivity). The skeleton was then represented as an undirected weighted graph using NetworkX, in which each skeleton voxel constitutes a node and edges connect adjacent voxels with weights equal to their Euclidean separation (1, √2, or √3 for face-, edge-, and corner-sharing neighbours, respectively). From this graph, the node-degree distribution, mean degree, maximum degree, number of edges, and the numbers of end points (degree = 1) and branch points (degree ≥ 3) were extracted. The network diameter was computed as the largest shortest-path length between nodes, restricted to the largest connected component for fragmented skeletons. Tortuosity was defined as the ratio of the longest geodesic (edge-weighted shortest) path between any pair of end points to the corresponding straight-line distance, T=L_geo/L_euc, where L_geo and L_euc denote the geodesic and Euclidean distances, respectively. Mitochondrial shape anisotropy was further quantified by principal component analysis (sklearn.decomposition.PCA) of the voxel coordinates, from which flatness (λ□/λ□) were derived from the ranked eigenvalues λ□≥ λ□≥ λ□.

#### Statistical analysis

Data normality was evaluated before applying statistical tests. Normally distributed datasets were analyzed using one-way ANOVA with Tukey’s post hoc multiple-comparisons test, while non-normally distributed datasets were analyzed using the Kruskal–Wallis test followed by Dunn’s post hoc test (*p < 0.05, **p < 0.01, ***p < 0.001, ****p < 0.0001). Statistical significance derived from ANOVA– Tukey analyses is indicated by black asterisks, whereas significance derived from Kruskal-Wallis– Dunn analyses is indicated by red asterisks. Brackets denote pairwise comparisons between treatment groups, and asterisks placed above whiskers denote comparisons between control and treatment groups. For the skeleton-derived graph parameters, mitochondrial morphotypes (fragmented, intermediate, and interconnected) were compared two groups at a time; accordingly, statistical significance was assessed using the two-sided Mann–Whitney U test for each pairwise comparison (statannotations), with significance indicated by asterisks above the corresponding box plots and Raincloud plot (*p < 0.05, **p < 0.01, ***p < 0.001, ****p < 0.0001).

## References

1. Westermann, B. (2010). Mitochondrial fusion and fission in cell life and death. Nat. Rev. Mol. Cell Biol. 11, 872–884. 10.1038/nrm3013.

2. Scott, I., and Youle, R.J. (2010). Mitochondrial fission and fusion. Essays Biochem. 47, 85. 10.1042/BSE0470085.

3. Lee, H., and Yoon, Y. (2014). Mitochondrial Fission: Regulation and ER Connection. Mol. Cells 37, 89–94. 10.14348/MOLCELLS.2014.2329.

4. Liu, Y., Mao, Z.H., Huang, J., Wang, H., Zhang, X., Zhou, X., Xu, Y., Pan, S., Liu, D., Liu, Z., et al. (2025). Mitochondria-Associated Endoplasmic Reticulum Membranes in Human Health and Diseases. MedComm (Beijing). 6, e70259. 10.1002/MCO2.70259.

5. Mookerjee, S.A., Goncalves, R.L.S., Gerencser, A.A., Nicholls, D.G., and Brand, M.D. (2015). The contributions of respiration and glycolysis to extracellular acid production. Biochimica et Biophysica Acta (BBA) - Bioenergetics 1847, 171–181. 10.1016/J.BBABIO.2014.10.005.

6. Chan, D.C. (2006). Mitochondrial fusion and fission in mammals. Annu. Rev. Cell Dev. Biol. 22, 79–99. 10.1146/ANNUREV.CELLBIO.22.010305.104638.

7. Chen, W., Zhao, H., and Li, Y. (2023). Mitochondrial dynamics in health and disease: mechanisms and potential targets. Signal Transduction and Targeted Therapy 2023 8:1 8, 333-. 10.1038/s41392-023-01547-9.

8. Jenkins, B.C., Neikirk, K., Katti, P., Claypool, S.M., Kirabo, A., McReynolds, M.R., and Hinton, A. (2024). Mitochondria in disease: changes in shapes and dynamics. Trends Biochem. Sci. 49, 346–360. 10.1016/J.TIBS.2024.01.011.

9. Zilocchi, M., Broderick, K., Phanse, S., Aly, K.A., and Babu, M. (2020). Mitochondria under the spotlight: On the implications of mitochondrial dysfunction and its connectivity to neuropsychiatric disorders. Comput. Struct. Biotechnol. J. 18, 2535–2546. 10.1016/J.CSBJ.2020.09.008.

10. Ong, S.B., Subrayan, S., Lim, S.Y., Yellon, D.M., Davidson, S.M., and Hausenloy, D.J. (2010). Inhibiting mitochondrial fission protects the heart against ischemia/reperfusion injury. Circulation 121, 2012–2022. 10.1161/CIRCULATIONAHA.109.906610/SUPPL_FILE/CIR200378VIDEO.AVI.

11. Wu, Q.R., Zheng, D.L., Liu, P.M., Yang, H., Li, L.A., Kuang, S.J., Lai, Y.Y., Rao, F., Xue, Y.M., Lin, J.J., et al. (2021). High glucose induces Drp1-mediated mitochondrial fission via the Orai1 calcium channel to participate in diabetic cardiomyocyte hypertrophy. Cell Death Dis. 12, 1–15. 10.1038/S41419-021-03502-4;TECHMETA.

12. Supale, S., Li, N., Brun, T., and Maechler, P. (2012). Mitochondrial dysfunction in pancreatic β cells. Trends in Endocrinology & Metabolism 23, 477–487. 10.1016/J.TEM.2012.06.002.

13. Haythorne, E., Rohm, M., van de Bunt, M., Brereton, M.F., Tarasov, A.I., Blacker, T.S., Sachse, G., Silva dos Santos, M., Terron Exposito, R., Davis, S., et al. (2019). Diabetes causes marked inhibition of mitochondrial metabolism in pancreatic β-cells. Nat. Commun. 10. 10.1038/s41467-019-10189-x.

14. Tseng, W.W., Chu, C.H., Lee, Y.J., Zhao, S., Chang, C., Ho, Y.P., and Wei, A.C. (2024). Metabolic regulation of mitochondrial morphologies in pancreatic beta cells: coupling of bioenergetics and mitochondrial dynamics. Commun. Biol. 7, 1267. 10.1038/s42003-024-06955-3.

15. Kim, J.A., Wei, Y., and Sowers, J.R. (2008). Role of Mitochondrial Dysfunction in Insulin Resistance. Circ. Res. 102, 401. 10.1161/CIRCRESAHA.107.165472.

16. Rodrigues, T., and Ferraz, L.S. (2020). Therapeutic potential of targeting mitochondrial dynamics in cancer. Biochem. Pharmacol. 182, 114282. 10.1016/J.BCP.2020.114282.

17. Li, A., Zhang, X., Singla, J., White, K., Loconte, V., Hu, C., Zhang, C., Li, S., Li, W., Francis, J.P., et al. (2022). Auto-segmentation and time-dependent systematic analysis of mesoscale cellular structure in β-cells during insulin secretion. PLoS One 17, e0265567. 10.1371/journal.pone.0265567.

18. Li, A., Zhang, S., Loconte, V., Liu, Y., Ekman, A., Thompson, G.J., Sali, A., Stevens, R.C., White, K., Singla, J., et al. (2022). An intensity-based post-processing tool for 3D instance segmentation of organelles in soft X-ray tomograms. PLoS One 17, e0269887. 10.1371/journal.pone.0269887.

19. Raveh, B., Sun, L., White, K.L., Sanyal, T., Tempkin, J., Zheng, D., Bharath, K., Singla, J., Wang, C., Zhao, J., et al. (2021). Bayesian metamodeling of complex biological systems across varying representations. Proc. Natl. Acad. Sci. U. S. A. 118, e2104559118. 10.1073/pnas.2104559118.

20. Singla, J., and White, K.L. (2021). A community approach to whole-cell modeling. Curr. Opin. Syst. Biol. 26, 33–38. 10.1016/J.COISB.2021.03.009.

21. Singla, J., McClary, K.M., White, K.L., Alber, F., Sali, A., and Stevens, R.C. (2018). Opportunities and Challenges in Building a Spatiotemporal Multi-scale Model of the Human Pancreatic β Cell. Cell 173, 11–19. 10.1016/j.cell.2018.03.014.

22. Leonard, A.P., Cameron, R.B., Speiser, J.L., Wolf, B.J., Peterson, Y.K., Schnellmann, R.G., Beeson, C.C., and Rohrer, B. (2015). Quantitative analysis of mitochondrial morphology and membrane potential in living cells using high-content imaging, machine learning, and morphological binning. Biochim. Biophys. Acta Mol. Cell Res. 1853, 348–360. 10.1016/j.bbamcr.2014.11.002.

23. Wiemerslage, L., and Lee, D. (2016). Quantification of mitochondrial morphology in neurites of dopaminergic neurons using multiple parameters. J. Neurosci. Methods 262, 56–65. 10.1016/j.jneumeth.2016.01.008.

24. Rambold, A.S., Kostelecky, B., Elia, N., and Lippincott-Schwartz, J. (2011). Tubular network formation protects mitochondria from autophagosomal degradation during nutrient starvation. Proc. Natl. Acad. Sci. U. S. A. 108, 10190–10195. 10.1073/pnas.1107402108.

25. Nikolaisen, J., Nilsson, L.I.H., Pettersen, I.K.N., Willems, P.H.G.M., Lorens, J.B., Koopman, W.J.H., and Tronstad, K.J. (2014). Automated quantification and integrative analysis of 2D and 3D mitochondrial shape and network properties. PLoS One 9. 10.1371/JOURNAL.PONE.0101365.

26. Chaudhry, A., Shi, R., and Luciani, D.S. (2020). A pipeline for multidimensional confocal analysis of mitochondrial morphology, function, and dynamics in pancreatic β-cells. Am. J. Physiol. Endocrinol. Metab. 318, E87–E101. 10.1152/AJPENDO.00457.2019.

27. Harwig, M.C., Viana, M.P., Egner, J.M., Harwig, J.J., Widlansky, M.E., Rafelski, S.M., and Hill, R.B. (2018). Methods for imaging mammalian mitochondrial morphology: a prospective on MitoGraph. Anal. Biochem. 552, 81. 10.1016/J.AB.2018.02.022.

28. Taguchi, K., Elias, B.C., Krystofiak, E., Qian, S., Sant, S., Yang, H., Fogo, A.B., and Brooks, C.R. (2021). Quantitative Super-Resolution Microscopy Reveals Promoting Mitochondrial Interconnectivity Protects against AKI. Kidney360 2, 1892–1907. 10.34067/KID.0001602021/-/DCSUPPLEMENTAL.

29. Ahmad, T., Aggarwal, K., Pattnaik, B., Mukherjee, S., Sethi, T., Tiwari, B.K., Kumar, M., Micheal, A., Mabalirajan, U., Ghosh, B., et al. (2013). Computational classification of mitochondrial shapes reflects stress and redox state. Cell Death Dis. 4. 10.1038/CDDIS.2012.213.

30. Song, W., Bossy, B., Martin, O.J., Hicks, A., Lubitz, S., Knott, A.B., and Bossy-Wetzel, E. (2008). Assessing mitochondrial morphology and dynamics using fluorescence wide-field microscopy and 3D image processing. Methods 46, 295–303. 10.1016/j.ymeth.2008.10.003.

31. Picard, M., Gentil, B.J., McManus, M.J., White, K., Louis, K.S., Gartside, S.E., Wallace, D.C., and Turnbull, D.M. (2013). Acute exercise remodels mitochondrial membrane interactions in mouse skeletal muscle. J. Appl. Physiol. 115, 1562. 10.1152/JAPPLPHYSIOL.00819.2013.

32. Ouellet, M., Guillebaud, G., Gervais, V., Lupien St-Pierre, D., and Germain, M. (2017). A novel algorithm identifies stress-induced alterations in mitochondrial connectivity and inner membrane structure from confocal images. PLoS Comput. Biol. 13, e1005612. 10.1371/JOURNAL.PCBI.1005612.

33. Rohani, A., Kashatus, J.A., Sessions, D.T., Sharmin, S., and Kashatus, D.F. (2020). Mito Hacker: a set of tools to enable high-throughput analysis of mitochondrial network morphology. Scientific Reports 2020 10:1 10, 18941-. 10.1038/s41598-020-75899-5.

34. Periasamy, A. (2013). Methods in cellular imaging.

35. Swift, L.H., and Colarusso, P. (2022). Fluorescence Microscopy: A Field Guide for Biologists. Methods in Molecular Biology 2440, 3–39. 10.1007/978-1-0716-2051-9_1.

36. Winey, M., Meehl, J.B., O’Toole, E.T., and Giddings, T.H. (2014). Conventional transmission electron microscopy. Mol. Biol. Cell 25, 319. 10.1091/MBC.E12-12-0863.

37. Weston, A.E., Armer, H.E.J., and Collinson, L.M. (2009). Towards native-state imaging in biological context in the electron microscope. J. Chem. Biol. 3, 101. 10.1007/S12154-009-0033-7.

38. Noske, A.B., Costin, A.J., Morgan, G.P., and Marsh, B.J. (2007). Expedited Approaches to Whole Cell Electron Tomography and Organelle Mark-Up in Situ in High-Pressure Frozen Pancreatic Islets. J. Struct. Biol. 161, 298. 10.1016/J.JSB.2007.09.015.

39. Denk, W., and Horstmann, H. (2004). Serial Block-Face Scanning Electron Microscopy to Reconstruct Three-Dimensional Tissue Nanostructure. PLoS Biol. 2, e329. 10.1371/JOURNAL.PBIO.0020329.

40. Narayan, K., and Subramaniam, S. (2015). Focused ion beams in biology. Nat. Methods 12, 1021–1031. 10.1038/NMETH.3623;SUBJMETA.

41. Lučić, V., Rigort, A., and Baumeister, W. (2013). Cryo-electron tomography: the challenge of doing structural biology in situ. J. Cell Biol. 202, 407–419. 10.1083/JCB.201304193.

42. Zhang, X., Carter, S.D., Singla, J., White, K.L., Butler, P.C., Stevens, R.C., and Jensen, G.J. (2020). Visualizing insulin vesicle neighborhoods in β cells by cryo-electron tomography. Sci. Adv. 6. 10.1126/sciadv.abc8258.

43. Gupta, A., Harrison, P.J., Wieslander, H., Pielawski, N., Kartasalo, K., Partel, G., Solorzano, L., Suveer, A., Klemm, A.H., Spjuth, O., et al. (2018). Deep Learning in Image Cytometry: A Review. Cytometry 95, 366. 10.1002/CYTO.A.23701.

44. Fogo, G.M., Anzell, A.R., Maheras, K.J., Raghunayakula, S., Wider, J.M., Emaus, K.J., Bryson, T.D., Bukowski, M.J., Neumar, R.W., Przyklenk, K., et al. (2021). Machine learning-based classification of mitochondrial morphology in primary neurons and brain. Sci. Rep. 11. 10.1038/s41598-021-84528-8.

45. Larabell, C.A., and Nugent, K.A. (2010). Imaging cellular architecture with X-rays. Curr. Opin. Struct. Biol. 20, 623–631. 10.1016/J.SBI.2010.08.008.

46. McDermott, G., Fox, D.M., Epperly, L., Wetzler, M., Barron, A.E., Le Gros, M.A., and Larabell, C.A. (2012). Visualizing and quantifying cell phenotype using soft X-ray tomography. BioEssays 34, 320–327. 10.1002/bies.201100125.

47. Do, M., Isaacson, S.A., McDermott, G., Le Gros, M.A., and Larabell, C.A. (2015). Imaging and characterizing cells using tomography. Arch. Biochem. Biophys. 581, 111. 10.1016/j.abb.2015.01.011.

48. Loconte, V., Singla, J., Li, A., Chen, J.H., Ekman, A., McDermott, G., Sali, A., Le Gros, M., White, K.L., and Larabell, C.A. (2022). Soft X-ray tomography to map and quantify organelle interactions at the mesoscale. Structure 30. 10.1016/j.str.2022.01.006.

49. Yadav, A., Singh, A., Deshmukh, A., Bharadwaj, P., Baliyan, A., White, K., and Singla, J. (2026). Robust mitochondria segmentation and morphological profiling using soft X-ray tomography. J. Struct. Biol. 218, 108291. 10.1016/J.JSB.2026.108291.

50. Loconte, V., and White, K.L. (2022). The use of soft X-ray tomography to explore mitochondrial structure and function. Mol. Metab. 57, 101421. 10.1016/J.MOLMET.2021.101421.

51. Deshmukh, A., Chang, K., Cuala, J., Hernandez Campos, M.J., Mahmood, S., Verma, R., Georgia, S., Loconte, V., and White, K.L. (2025). Secretory stimuli distinctly regulate insulin secretory granule maturation through structural remodeling. Structure 33, 1831-1843.e4. 10.1016/j.str.2025.07.022.

52. White, K.L., Singla, J., Loconte, V., Chen, J.H., Ekman, A., Sun, L., Zhang, X., Francis, J.P., Li, A., Lin, W., et al. (2020). Visualizing subcellular rearrangements in intact β cells using soft x-ray tomography. Sci. Adv. 6. 10.1126/sciadv.abc8262.

53. Lounas, A., Lebrun, A., Laflamme, I., Vernoux, N., Savage, J., Tremblay, M.È., Germain, M., and Richard, F.J. (2022). A 3D analysis revealed complexe mitochondria morphologies in porcine cumulus cells. Scientific Reports 2022 12:1 12, 15403-. 10.1038/s41598-022-19723-2.

54. Brenner, B., Xu, F., Zhang, Y., Kweon, J., Fang, R., Sheibani, N., Zhang, S.X., Sun, C., and Zhang, H.F. (2024). Quantifying nanoscopic alterations associated with mitochondrial dysfunction using three-dimensional single-molecule localization microscopy. Biomed. Opt. Express 15, 1571. 10.1364/BOE.510351.

55. Zahedi, A., On, V., Phandthong, R., Chaili, A., Remark, G., Bhanu, B., and Talbot, P. Deep Analysis of Mitochondria and Cell Health Using Machine Learning. 10.1038/s41598-018-34455-y.

56. Jia, X., Mao, D., Guo, J., Ke, J., Zhu, Y., Zhao, X., Luo, Z., Liu, X., Tang, R., Hou, R., et al. (2024). Epigallocatechin gallate attenuated high glucose-induced pancreatic beta cell dysfunction by modulating DRP1-mediated mitochondrial apoptosis pathways. Scientific Reports 2024 14:1 14, 16809-. 10.1038/s41598-024-67867-0.

57. Jhun, B.S., Lee, H., Jin, Z.G., and Yoon, Y. (2013). Glucose Stimulation Induces Dynamic Change of Mitochondrial Morphology to Promote Insulin Secretion in the Insulinoma Cell Line INS-1E. PLoS One 8. 10.1371/journal.pone.0060810.

58. Pandey, S., Mangmool, S., Madreiter-Sokolowski, C.T., Wichaiyo, S., Luangmonkong, T., and Parichatikanond, W. (2023). Exendin-4 protects against high glucose-induced mitochondrial dysfunction and oxidative stress in SH-SY5Y neuroblastoma cells through GLP-1 receptor/Epac/Akt signaling. Eur. J. Pharmacol. 954, 175896. 10.1016/J.EJPHAR.2023.175896.

59. Al Sabaani, N. (2021). Exendin-4 inhibits high glucose-induced oxidative stress in retinal pigment epithelial cells by modulating the expression and activation of p66Shc. Cutan. Ocul. Toxicol. 40, 175–186. 10.1080/15569527.2020.1844727.

60. Khine, H.E.E., Mangmool, S., and Parichatikanond, W. (2025). Exendin-4 improves mitochondrial integrity against cisplatin-induced cardiac damage: Targeting p53 and NF-κB pathways. Eur. J. Pharmacol. 1007, 178233. 10.1016/J.EJPHAR.2025.178233.

61. Bereiter-Hahn, J. (1990). Behavior of Mitochondria in the Living Cell. Int. Rev. Cytol. 122, 1–63. 10.1016/S0074-7696(08)61205-X.

62. Rafelski, S.M. (2013). Mitochondrial network morphology: Building an integrative, geometrical view. BMC Biol. 11. 10.1186/1741-7007-11-71.

63. Li, Z., Zhou, Z., Huang, G., Hu, F., Xiang, Y., and He, L. (2013). Exendin-4 Protects Mitochondria from Reactive Oxygen Species Induced Apoptosis in Pancreatic Beta Cells. PLoS One 8, e76172. 10.1371/JOURNAL.PONE.0076172.

64. Wei, Q., Sun, Y.Q., and Zhang, J. (2012). Exendin-4, a glucagon-like peptide-1 receptor agonist, inhibits cell apoptosis induced by lipotoxicity in pancreatic β-cell line. Peptides (N.Y.). 37, 18–24. 10.1016/j.peptides.2012.06.018.

65. Youle, R.J., and Van Der Bliek, A.M. (2012). Mitochondrial fission, fusion, and stress. Science (1979). 337, 1062–1065. 10.1126/SCIENCE.1219855.

66. Tondera, D., Grandemange, S., Jourdain, A., Karbowski, M., Mattenberger, Y., Herzig, S., Da Cruz, S., Clerc, P., Raschke, I., Merkwirth, C., et al. (2009). SlP-2 is required for stress-induced mitochondrial hyperfusion. EMBO Journal 28, 1589–1600. 10.1038/emboj.2009.89.

67. Liesa, M., and Shirihai, O.S. (2013). Mitochondrial dynamics in the regulation of nutrient utilization and energy expenditure. Cell Metab. 17, 491–506. 10.1016/j.cmet.2013.03.002.

68. Ekman, A.A., Chen, J.H., Guo, J., McDermott, G., Le Gros, M.A., and Larabell, C.A. (2017). Mesoscale imaging with cryo-light and X-rays: Larger than molecular machines, smaller than a cell. Biol. Cell 109, 24–38. 10.1111/boc.201600044.

69. Egan, J.M., Clocquet, A.R., and Elahi, D. (2002). The Insulinotropic Effect of Acute Exendin-4 Administered to Humans: Comparison of Nondiabetic State to Type 2 Diabetes. J. Clin. Endocrinol. Metab. 87, 1282–1290. 10.1210/jcem.87.3.8337.

70. Kang, M.Y., Oh, T.J., and Cho, Y.M. (2015). Glucagon-Like Peptide-1 Increases Mitochondrial Biogenesis and Function in INS-1 Rat Insulinoma Cells. Endocrinology and Metabolism 30, 216. 10.3803/ENM.2015.30.2.216.

71. Wai, T., and Langer, T. (2016). Mitochondrial Dynamics and Metabolic Regulation. Trends in Endocrinology & Metabolism 27, 105–117. 10.1016/J.TEM.2015.12.001.

72. Cogliati, S., Enriquez, J.A., and Scorrano, L. (2016). Mitochondrial Cristae: Where Beauty Meets Functionality. Trends Biochem. Sci. 41, 261–273. 10.1016/j.tibs.2016.01.001.

73. Glancy, B. (2020). Visualizing Mitochondrial Form and Function within the Cell. Trends Mol. Med. 26, 58–70. 10.1016/j.molmed.2019.09.009.

74. Toyama, E.Q., Herzig, S., Courchet, J., Lewis, T.L., Losón, O.C., Hellberg, K., Young, N.P., Chen, H., Polleux, F., Chan, D.C., et al. (2016). Metabolism. AMP-activated protein kinase mediates mitochondrial fission in response to energy stress. Science 351, 275–281. 10.1126/science.aab4138.

75. Mishra, P., and Chan, D.C. (2016). Metabolic regulation of mitochondrial dynamics. Journal of Cell Biology 212, 379–387. 10.1083/JCB.201511036.

76. Ding, Y., Li, J., Zhang, J., Li, P., Bai, H., Fang, B., Fang, H., Huang, K., Wang, G., Nowell, C.J., et al. (2025). Mitochondrial segmentation and function prediction in live-cell images with deep learning. Nature Communications 2025 16:1 16, 1–15. 10.1038/s41467-025-55825-x.

77. Perez, L., Ng, X.W., Mohs, M., Piston, D.W., and Mukherji, S. (2025). Mitochondrial position responds to glucose stimulation in a model of the pancreatic beta cell. Biophys. J. 125, 168. 10.1016/j.bpj.2025.11.018.

78. Parichatikanond, W., Pandey, S., and Mangmool, S. (2024). Exendin-4 exhibits cardioprotective effects against high glucose-induced mitochondrial abnormalities: Potential role of GLP-1 receptor and mTOR signaling. Biochem. Pharmacol. 229, 116552. 10.1016/j.bcp.2024.116552.

79. Huang, C., Deng, K., and Wu, M. (2023). Mitochondrial cristae in health and disease. Int. J. Biol. Macromol. 235, 123755. 10.1016/j.ijbiomac.2023.123755.

80. Serikbaeva, A., Li, Y., Ganesh, B., Zelkha, R., and Kazlauskas, A. (2022). Hyperglycemia Promotes Mitophagy and Thereby Mitigates Hyperglycemia-Induced Damage. Am. J. Pathol. 192, 1779–1794. 10.1016/J.AJPATH.2022.08.004.

81. Oh, S.J., Park, K., Sonn, S.K., Oh, G.T., and Lee, M.S. (2023). Pancreatic β-cell mitophagy as an adaptive response to metabolic stress and the underlying mechanism that involves lysosomal Ca2+ release. Exp. Mol. Med. 55, 1922–1932. 10.1038/s12276-023-01055-4.

82. Schindelin, J., Arganda-Carreras, I., Frise, E., Kaynig, V., Longair, M., Pietzsch, T., Preibisch, S., Rueden, C., Saalfeld, S., Schmid, B., et al. (2012). Fiji: an open-source platform for biological-image analysis. Nature Methods 2012 9:7 9, 676–682. 10.1038/nmeth.2019.

83. Kremer, J.R., Mastronarde, D.N., and McIntosh, J.R. (1996). Computer Visualization of Three-Dimensional Image Data Using IMOD. J. Struct. Biol. 116, 71–76. 10.1006/jsbi.1996.0013.

84. Meng, E.C., Goddard, T.D., Pettersen, E.F., Couch, G.S., Pearson, Z.J., Morris, J.H., and Ferrin, T.E. (2023). UCSF ChimeraX: Tools for structure building and analysis. Protein Sci. 32. 10.1002/pro.4792.

85. Le Gros, M.A., Mcdermott, G., Cinquin, B.P., Smith, E.A., Do, M., Chao, W.L., Naulleau, P.P., and Larabell, C.A. (2014). Biological soft X-ray tomography on beamline 2.1 at the Advanced Light Source. J. Synchrotron Radiat. 21, 1370–1377. 10.1107/S1600577514015033.

86. Parkinson, D.Y., Knoechel, C., Yang, C., Larabell, C.A., and Le Gros, M.A. (2012). Automatic alignment and reconstruction of images for soft X-ray tomography. J. Struct. Biol. 177, 259–266. 10.1016/j.jsb.2011.11.027.

87. Koopman, W.J.H., Visch, H.-J., Verkaart, S., Van Den Heuvel, L.W.P.J., Smeitink, J.A.M., Willems, P.H.G.M., and Van Den Heuvel, L.-B.W.P.J. (2005). Mitochondrial network complexity and pathological decrease in complex I activity are tightly correlated in isolated human complex I deficiency. Am J Physiol Cell Physiol 289, 881–890. 10.1152/ajpcell.00104.2005.-Com.

88. Vincent, A.E., White, K., Davey, T., Philips, J., Ogden, R.T., Lawess, C., Warren, C., Hall, M.G., Ng, Y.S., Falkous, G., et al. (2019). Quantitative 3D Mapping of the Human Skeletal Muscle Mitochondrial Network. Cell Rep. 26, 996–1009.e4. 10.1016/j.celrep.2019.01.010.

89. Blender - The Free and Open Source 3D Creation Software — blender.org https://www.blender.org/.

90. Chang, K., Deshmukh, A., Verma, R., Loconte, V., and White, K.L. (2025). Classifying Biophysical Subpopulations of Insulin Secretory Granules using Quantitative Whole Cell Structure Analysis. bioRxiv. 10.1101/2025.09.09.675239.

