## Supplementary Tables and Figures for "Morphotype-Resolved 3D Morphometry Reveals a Structure–Density–Location Coupling in Mitochondrial Networks" (from another version of this preprint: version 1, 2026-03-20)

**Table S1**. **Morphological features used for mitochondrial clustering analysis:** This table lists the quantitative morphological features extracted from 3D mitochondrial reconstructions that were used for clustering analysis. These descriptors capture mitochondrial size, shape, and structural complexity, enabling classification of mitochondria into distinct morphological clusters.

| Feature Name | Description |
| --- | --- |
| Volume  (µm^3^) | The actual physical 3D space the mitochondrion occupies.  **Used for:** Distinguishing between tiny, fragmented mitochondria and massive, fused mitochondrial networks. |
| Convex hull volume (µm^3^) | The volume of the tightest possible invisible boundary wrapping around the entire mitochondrion (like shrink-wrap).  **Used for:** Detecting complex or branched structures. If a mitochondrion is highly branched or U-shaped, its "shrink-wrap" volume will be much larger than its actual volume. |
| Sphericity | A score of how perfectly round the structure is (where 1.0 is a perfect sphere).  **Used for:** Identifying swollen or severely fragmented mitochondria, which tend to form perfect little spheres rather than healthy elongated tubes. |
| Aspect ratio | The proportion of the mitochondrion's longest dimension to its shortest. **Used for:** Quantifying how "stretched" or thread-like the structure is. High values indicate long, tubular mitochondria. |
| Flatness | A measure of how "squashed" or pancake-like the mitochondrion is.  **Used for:** Differentiating standard cylindrical mitochondrial tubes from abnormal, sheet-like structures that can form under certain cellular conditions. |
| Skeleton length | The total length of the mitochondrion's central "spine," ignoring how thick or thin the outer walls are.  **Used for:** Measuring the true reach and span of a mitochondrial network, regardless of whether it is swollen or thin. |
| Tortuosity | A measure of how twisted, curled, or winding the mitochondrion is compared to a perfectly straight line.  **Used for:** Identifying highly folded or zigzagging structures, which can indicate cellular stress, spatial crowding, or specific mobility states. |
| Fractal dimension | A mathematical measure of how incredibly complex or highly textured the structure is.  **Used for:** Quantifying extreme intricacy, such as highly ruffled surfaces or densely packed, chaotic branching that standard shape metrics can't capture. |

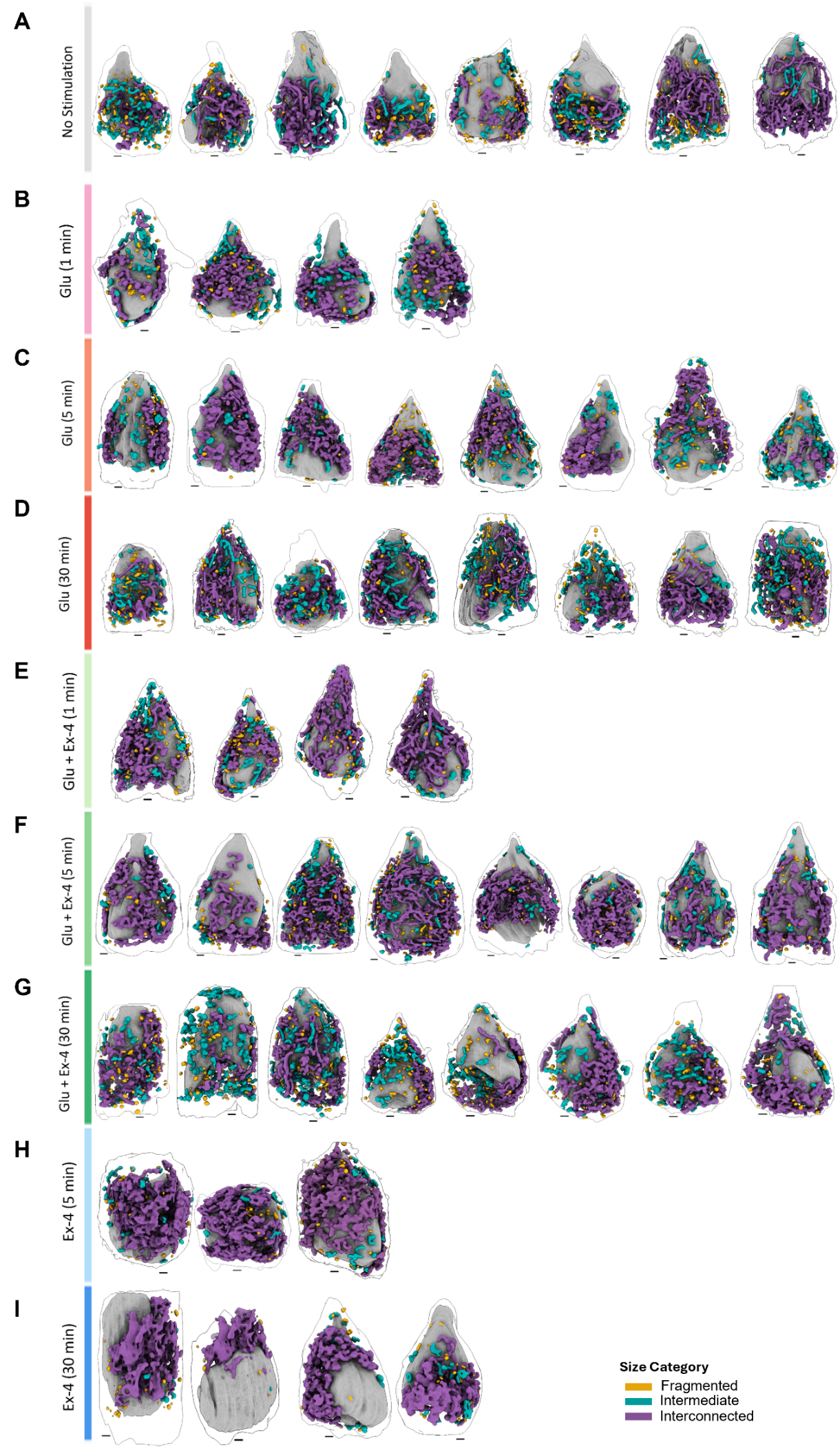

**Figure S1**. **3D illustrations of INS-1E cells under different stimulation conditions**

Three-dimensional renderings of whole INS-1E cells generated from segmented organelle masks derived from soft X-ray tomography (SXT) datasets: (A) No Stimulation, (B) Glucose (1 min), (C) Glucose (5 min), (D) Glucose (30 min), (E) Glucose + Ex-4 (1 min), (F) Glucose + Ex-4 (5 min), (G) Glucose + Ex-4 (30 min), (H) Ex-4 (5 min) and (I) Ex-4 (30 min).

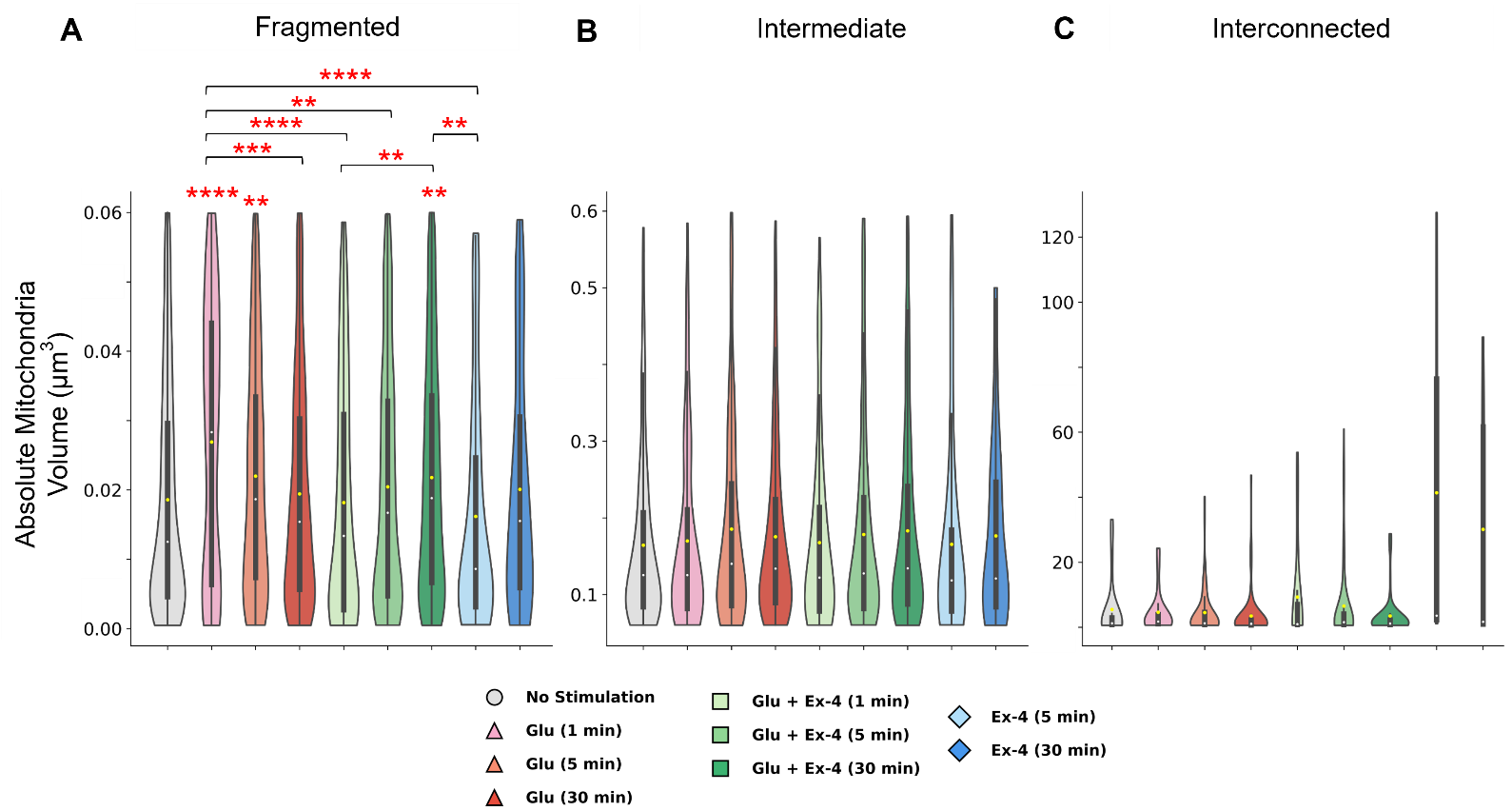

**Figure S2.** **Absolute mitochondrial volume across morphotypes and stimulation conditions**

Violin plots showing the distribution of absolute mitochondrial volume (µm³), related to Figure 2A, for individual mitochondria classified as (A) Fragmented, (B) Intermediate, and (C) Interconnected morphotypes across experimental conditions. The yellow dot indicates the mean, and the white dot indicates the median. Statistical significance relative to the unstimulated (NS) control is shown by asterisks above individual conditions, while brackets denote significant pairwise differences between stimulated groups. Red asterisks represent Kruskal–Wallis tests followed by Dunn’s post hoc comparisons (*p < 0.05, **p < 0.01, ***p < 0.001, ****p < 0.0001).

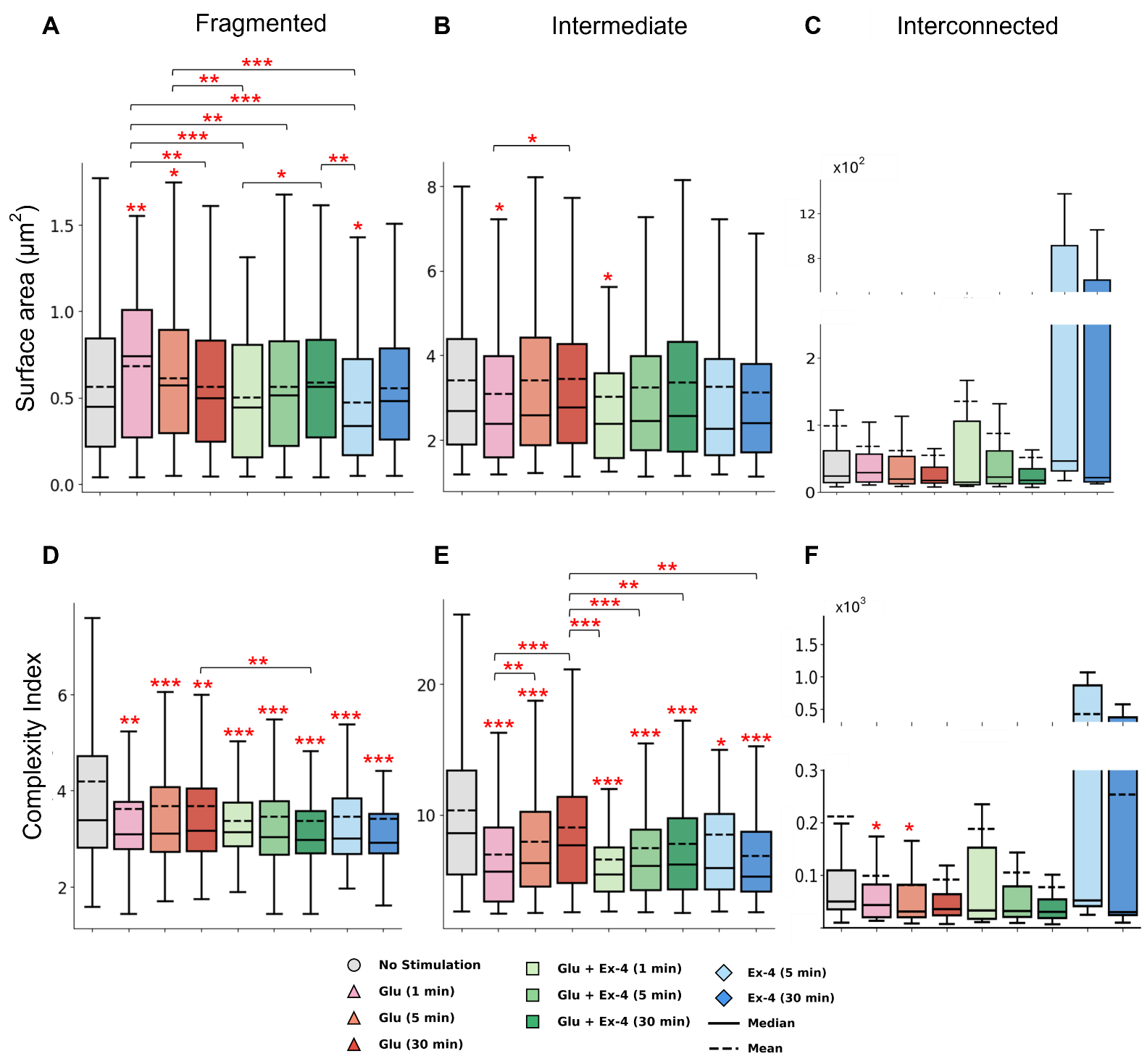

**Figure S3. Mitochondrial surface area and complexity index across morphotypes**

Box plots showing the distribution of mitochondrial surface area (µm²) (A-C) and complexity index (D-F), related to Figure 2B-C, Surface area and complexity index were computed from segmented SXT-derived 3D masks. Solid lines represent medians and dashed lines represent means.

Asterisks above individual conditions indicate statistical significance relative to the unstimulated (NS) control, while brackets denote significant pairwise differences between stimulated groups. Red asterisks represent Kruskal–Wallis tests followed by Dunn’s post hoc comparisons (*p < 0.05, **p < 0.01, ***p < 0.001, ****p < 0.0001).

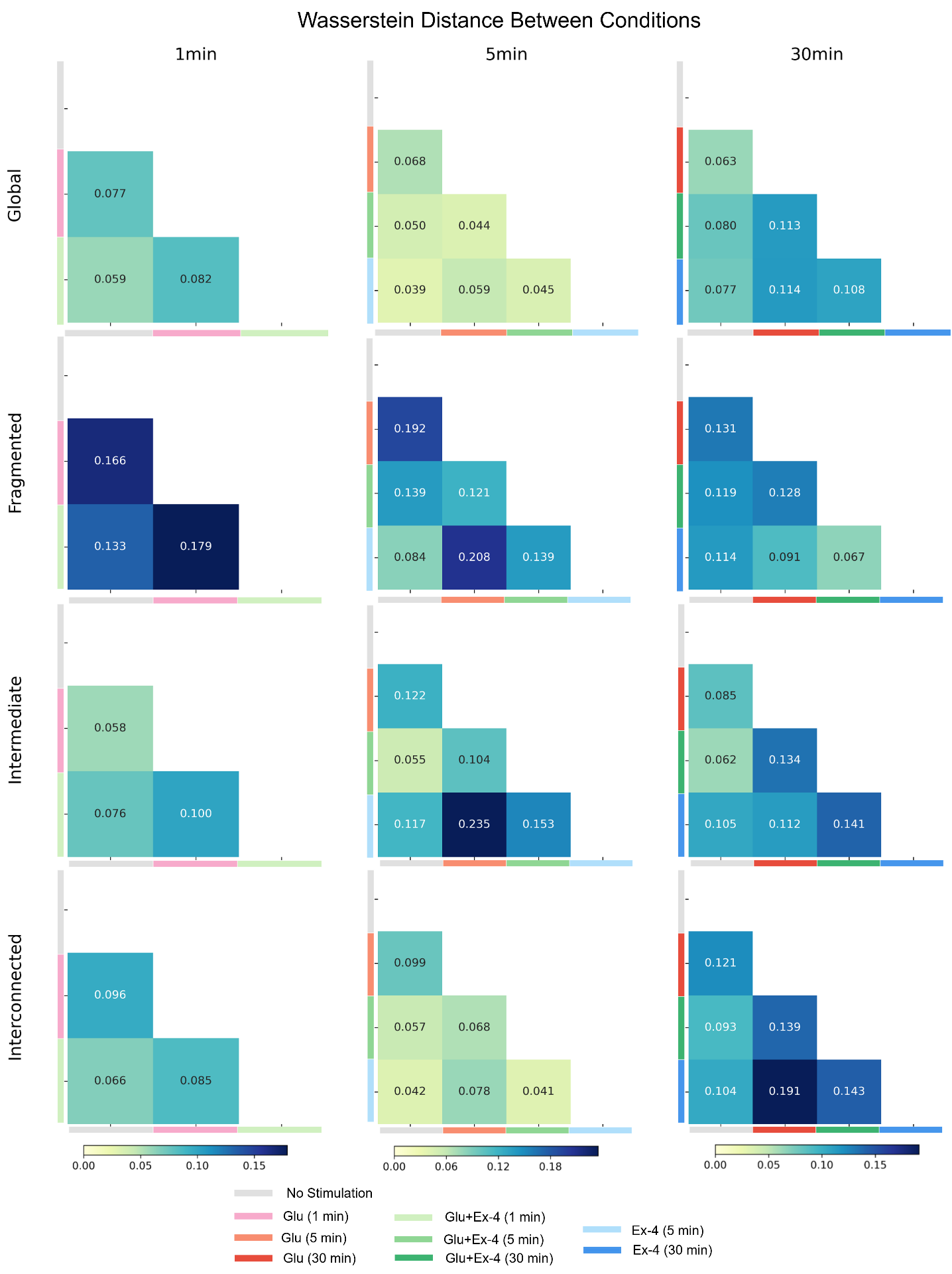

**Figure S4. Pairwise comparison of mitochondrial spatial distributions using Wasserstein distance**
(A–D) Heatmaps showing pairwise Wasserstein distances between stimulation conditions, computed from morphotype-specific spatial distributions. Each cell represents the distributional dissimilarity between two conditions. Representative 3D renderings illustrate the corresponding morphotype analyzed in each panel. Color bars denote Wasserstein distance magnitude.

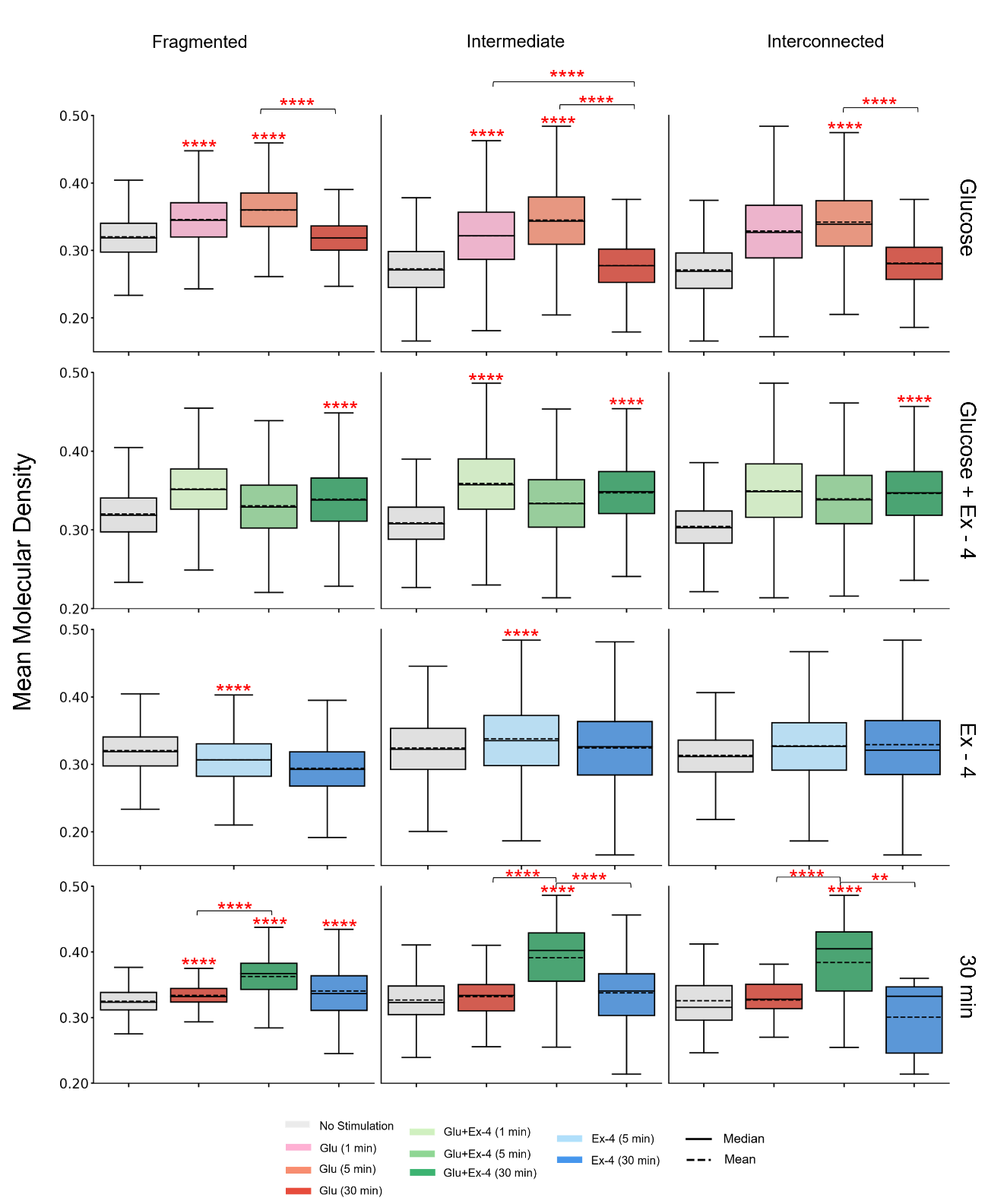

**Figure S5. Mean mitochondrial molecular density across stimulation conditions and morphotypes**

Box plots showing the distribution of mean mitochondrial molecular density across experimental conditions, related to Figure 3. (A) Fragmented, (B)Intermediate, (C) Interconnected, treatment including No Stimulation (NS), Glucose (1, 5, 30 min), Glucose + Ex-4 (1, 5, 30 min), and Ex-4 (5, 30 min). Solid lines represent medians and dashed lines represent means.

Red asterisks above individual conditions indicate statistical significance relative to the unstimulated (NS) control, while brackets denote significant pairwise comparisons between stimulated groups. Red asterisks correspond to Kruskal–Wallis tests followed by Dunn’s post hoc multiple-comparison analysis (*p < 0.05, **p < 0.01, ***p < 0.001, ****p < 0.0001).
